# Intraperitoneal activation of myeloid cells clears ascites and reveals IL27-dependent regression of metastatic ovarian cancer

**DOI:** 10.1101/2024.06.25.600597

**Authors:** Brennah Murphy, Taito Miyamoto, Bryan S. Manning, Gauri Mirji, Alessio Ugolini, Toshitha Kannan, Kohei Hamada, Yanfang Peipei Zhu, Daniel T. Claiborne, Lu Huang, Rugang Zhang, Yulia Nefedova, Andrew Kossenkov, Filippo Veglia, Rahul Shinde, Nan Zhang

## Abstract

Patients with metastatic ovarian cancer (OvCa) have a 5-year survival rate of less than 30% due to persisting dissemination of chemoresistant cells in the peritoneal fluid and the immunosuppressive microenvironment in the peritoneal cavity. Here, we report that intraperitoneal administration of β-glucan and IFNγ (BI) induced robust tumor regression in clinically relevant models of metastatic OvCa. BI induced tumor regression by controlling fluid tumor burden and activating localized antitumor immunity. β-glucan alone cleared ascites and eliminated fluid tumor cells by inducing intraperitoneal clotting in the fluid and Dectin-1-Syk-dependent NETosis in the omentum. In omentum tumors, BI expanded a novel subset of immunostimulatory IL27+ macrophages and neutralizing IL27 impaired BI efficacy *in vivo*. Moreover, BI directly induced IL27 secretion in macrophages where single agent treatment did not. Finally, BI extended mouse survival in a chemoresistant model and significantly improved chemotherapy response in a chemo-sensitive model. In summary, we propose a new therapeutic strategy for the treatment of metastatic OvCa.

## Introduction

Ovarian cancer (OvCa) is the most lethal gynecological cancer in the United States and the fifth leading cause of cancer-related deaths in women due in large part to metastases (Siegel et al., 2023). Unlike other cancers that metastasize through circulation, OvCa cells mostly disseminate directly into the peritoneal cavity and preferentially seed in the omentum prior to metastasizing to other peritoneal organs (Ma, 2020). Because it is difficult to detect at early stages, OvCa is often diagnosed at later stages with metastatic lesions present throughout the peritoneal cavity (stage III and IV) (Lengyel, 2010). Despite an initial positive response to chemotherapy, most patients relapse and ultimately present with intraperitoneal chemoresistant disease and poor prognosis (Colombo et al., 2017). Meanwhile, recent breakthroughs in immune checkpoint therapies have led to little improvement in OvCa prognosis (Monk et al., 2021; Pujade-Lauraine et al., 2021). Therefore, there is an urgent need to develop alternative therapies for end-stage metastatic OvCa. Multiple mechanisms contribute to therapy resistance in metastatic OvCa. First, the presence of disseminated cancer cells in the peritoneal fluid following initial treatment may promote therapy resistance in relapsed patients (Shield et al., 2009). A growing body of evidence suggests that these disseminating cells, which are present in malignant ascites and cannot be surgically resected, exhibit cancer stem cell characteristics that render them highly invasive and broadly resistant to therapy (Latifi et al., 2012; Liao et al., 2014; Shepherd and Dick, 2022; Shield et al., 2009; Ward Rashidi et al., 2019). Targeting these cells remains a significant challenge to overcome therapy resistance and relapse.

The second mechanism of therapy resistance is likely driven by the highly immunosuppressive microenvironment in the peritoneal cavity (Almeida-Nunes et al., 2022). Its immunosuppressive nature is predominantly supported by myeloid cells, namely macrophages (MΦs) and neutrophils, which are the most abundant cell type found in OvCa tumors and malignant ascites (Charoentong et al., 2017; Izar et al., 2020; Lee et al., 2019; Raghavan et al., 2019; Rickard et al., 2021). The peritoneal cavity consists of at least three anatomical compartments: the peritoneal fluid, the omentum, and the peritoneal membrane. Recent studies have revealed that tissue-resident MΦs in all three compartments can suppress immune responses and promote OvCa progression (Casanova-Acebes et al., 2020; Etzerodt et al., 2020; Long et al., 2021; Miyamoto et al., 2023; Zhang et al., 2021a). Of note, omental MΦs are known to contribute to the formation the pre-metastatic niche (Etzerodt 2020, Krishan 2020) and neutrophils recruited to the omentum during early OvCa progression have also been reported to promote metastasis into the omentum (Lee et al., 2019). Therefore, alternative immunotherapy approaches directly targeting myeloid cells in the peritoneal cavity may be necessary to overcome disease progression and resistance.

Nearly a century ago, it was discovered that administration of dead pathogens (specifically Coley’s toxin) could stimulate an antitumor response in some patients, likely via activating myeloid cells that promoted cancer killing (Wiemann and Starnes, 1994). Here, we sought to utilize a similar strategy to treat metastatic OvCa by administering β-glucan alongside interferon gamma (IFNγ) to activate myeloid cells in the peritoneal cavity. β-glucan is a polysaccharide derived from yeast cell walls and a known activator of myeloid cells. It has been shown to inhibit tumor progression in several non-OvCa tumor models (Bradner et al., 1958; Cheung et al., 2002; Hong et al., 2004; Kalafati et al., 2020; Woeste et al., 2023). Whether it can inhibit metastatic OvCa is not known. IFNγ is an immunogenic cytokine that is essential for innate and adaptive immunity. It was originally identified as a MΦ activation factor (Celada et al., 1984; Schreiber et al., 1982; Svedersky et al., 1984) and is crucial for reprogramming tumor associated MΦs (TAMs) in multiple tumor models (Alspach et al., 2019; Sun et al., 2021). Importantly, IFNγ alone failed in the most recent clinical trial to improve prognosis of patients with metastatic OvCa, indicating additional activation signals are necessary (Miller et al., 2009). Both β-glucan and IFNγ are currently in separate clinical trials to investigate efficacy in treating multiple cancers (NCT05159778 and NCT04628338). Importantly, emerging evidence suggests that therapies combining IFNγ and pathogen-derived molecules (e.g., β-glucan or lipopolysaccharide [LPS]) can reverse the immunosuppressive microenvironments in a few clinically relevant, orthotopic cancer models (Sun et al., 2021; Wattenberg et al., 2023). However, whether or how these therapies could treat metastatic OvCa remains poorly understood.

Here, we report that the intraperitoneal administration of β-glucan and IFNγ (BI) successfully induced the robust regression of metastatic OvCa tumors and controlled cancer metastasis. In the peritoneal fluid, we identified two complementary mechanisms through which β-glucan eliminates cancer cells: one that requires macrophage-mediated clotting in the peritoneal fluid and another that requires Dectin-1-Syk-dependent NETosis in the omentum. In solid metastases, we found that both agents for BI treatment are required for tumor regression. BI induced anti-tumor immunity in omentum tumors in part via MΦ-derived IL27. *In vitro*, BI directly induced IL27 secretion in MΦs which consequently activates CD8^+^ T cells. Moreover, we found that higher IL27 expression predicted better overall survival in patients with metastatic OvCa. Overall, this study proposes a new promising therapeutic approach for treating metastatic OvCa and reveals novel mechanisms of tumor control in the peritoneal fluid and omentum tumors.

## Results

### β-glucan significantly reduces ovarian cancer fluid tumor burden

As β-glucan is a well-known activator of innate immunity and has been reported to control other tumor types *in vivo* (Bradner et al., 1958; Cheung et al., 2002; Hong et al., 2004; Kalafati et al., 2020; Woeste et al., 2023), we first evaluated whether β-glucan alone could effectively treat tumors in the commonly utilized ID8 mouse model of OvCa. Luciferase- and GFP-tagged ID8 cells were seeded intraperitoneally (i.p.) to simulate metastatic OvCa and mice were treated once every two weeks with β-glucan (Figure S1A). Mouse tumor burden was significantly decreased in β-glucan treated mice (Figure 1A). Additionally, malignant ascites (the accumulation of tumor cell- and red blood cell-containing fluid in the peritoneal cavity – a hallmark of metastatic disease) was completely inhibited (Figure 1B) and ID8 cells were undetectable in the peritoneal lavage following β-glucan treatment (Figure 1C). This data indicates that β-glucan is sufficient to control ID8 tumor progression *in vivo*.

**Figure 1.**
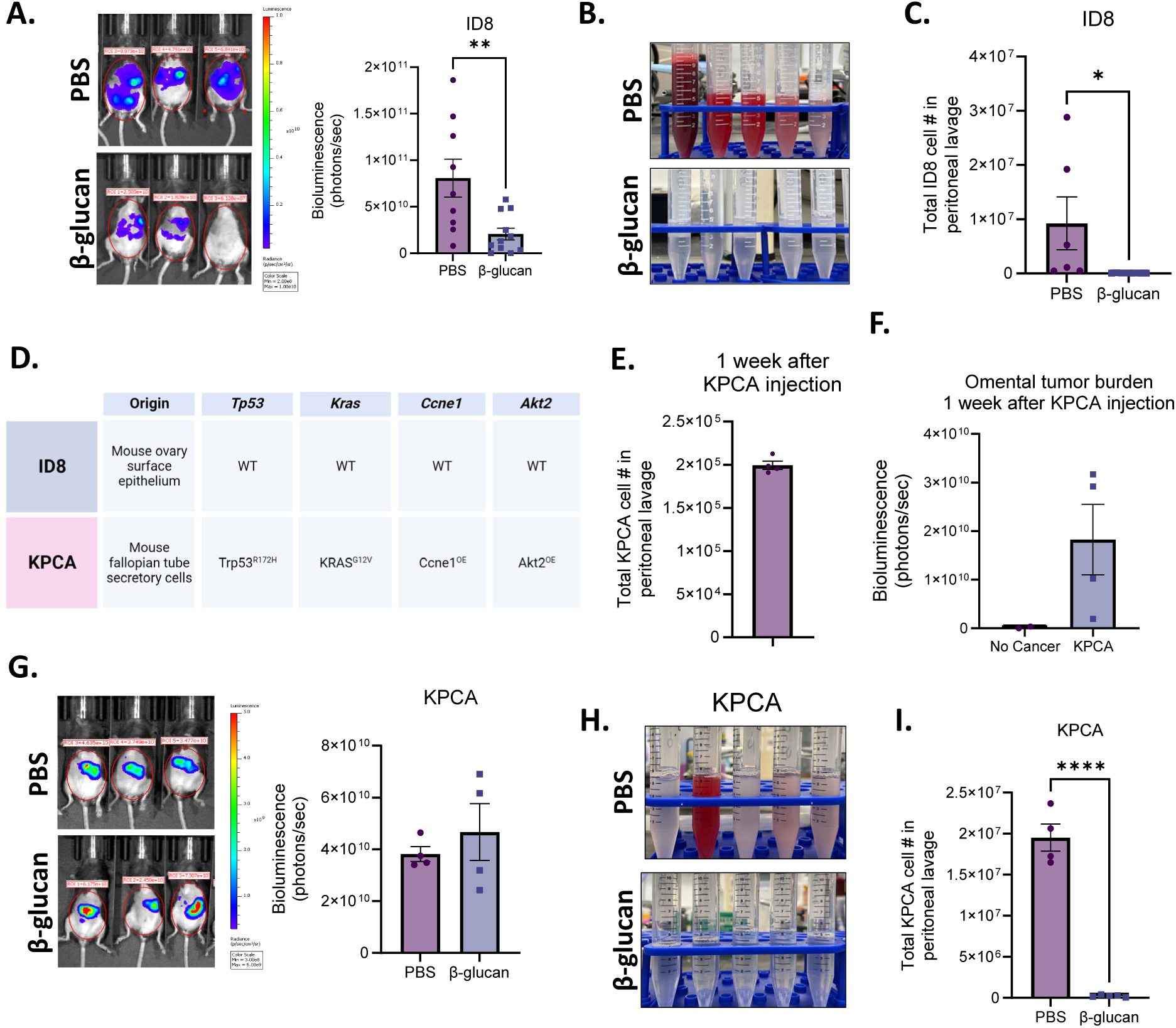
β-glucan significantly reduces OvCa fluid tumor burden. (A) Representative bioluminescence images and quantification of bioluminescence signals in PBS- and β-glucan-treated ID8 tumor-bearing mice 42 days post tumor-seeding. (B) Representative pictures of peritoneal lavage and (C) quantification of GFP^+^ ID8 OvCa cells in the peritoneal lavage of PBS- and β-glucan-treated mice. (D) Tissue of origin and mutation status of *Trp53, Kras, Ccne1,* and *Akt2* in ID8 and KPCA OvCa cell lines. (E) Quantification of KPCA cells in the peritoneal lavage one week after KPCA seeding. (F) quantification of omentum bioluminescence signals one week after KPCA seeding. (G) Representative bioluminescence images and quantification of bioluminescence signals in PBS- and β-glucan-treated KPCA tumor-bearing mice. (H) Representative pictures of peritoneal lavage and (I) quantification of GFP^+^ KPCA OvCa cells in the peritoneal lavage in PBS- and β-glucan-treated mice. Student’s t test was used *p<0.05 ; ****p<0.0001. Error bars are standard errors of the mean.

Although widely utilized for the past 20 years, it is now appreciated that ID8 cells do not harbor any common patient-relevant mutations or somatic copy-number alterations seen in human OvCa (Iyer et al., 2021; Roby et al., 2000; Walton et al., 2016). Therefore, we next wanted to test β-glucan efficacy in another syngeneic model of OvCa that utilizes a recently characterized cell line containing OvCa patient-relevant genetic alterations, KPCA (*KRAS*^G12V^*Trp53*^R172H^*Ccne1*^Overexpression (OE)^*Akt2*^OE^, Figure 1D) (Iyer et al., 2021). *KRAS, TP53, CCNE1,* and *AKT2* are mutant alleles observed in high grade serous ovarian tumors with frequencies of 12%, 96%, 19%, and 6% respectively (Iyer et al., 2021; Zhang et al., 2021b). Interestingly, amplification of *KRAS* and *CCNE1* was recently identified as a marker for chemoresistant OvCa (Smith et al., 2023) and these tumors were indeed reported to be resistant to chemotherapy (Iyer et al., 2021). Therefore, utilizing this model is valuable for developing effective therapies against therapy resistant OvCa in patients, an unmet clinical need.

Mice bearing KPCA tumors have a median survival of only 35 days, which is a nearly 3-fold decrease compared to the ID8 model which has a median survival of 114 days (Iyer et al., 2021; Roby et al., 2000). To account for this, the experiment design was modified to initiate the treatment 1 week following i.p. seeding of KPCA cells, and mice were treated with two doses of β-glucan on days 7 and 14 (Figure S1B). We first confirmed the presence of KPCA cells (GFP^+^luciferase^+^) in the peritoneal fluid (Figure 1E & S1C) as well as in the omentum at the time of treatment initiation (Figure 1F, S1D & S1E), which models metastatic OvCa (stage III or IV) in patients (Prat, 2014). For reasons that are not known, KPCA cells lose their luminescence in the peritoneal fluid. Therefore, bioluminescent signal is indicative of solid tumor burden, and fluid tumor burden is exclusively analyzed by flow cytometric analysis by GFP+ signal. In contrast to ID8 tumors, β-glucan could not reduce KPCA solid tumor burden (Figure 1G). However, β-glucan once again was able to inhibit accumulation of malignant ascites (Figure 1H & S1F) and significantly reduced KPCA presence in the peritoneal lavage (Figure 1I). Taken together this data suggests that β-glucan is sufficient to control fluid tumor burden independent of tumor type. However, β-glucan was unable to control metastatic growth of KPCA tumors. The ability of β-glucan to control solid tumor progression of ID8 tumors but not KPCA further highlights the critical role of tumor mutation status on therapy response and emphasizes the importance of utilizing preclinical models which better model human cancer.

### β-glucan captures ovarian cancer into solid nodular structures via intraperitoneal clotting and Dectin-1-Syk-dependent NETosis in the omentum

Given that β-glucan significantly reduced the presence of cancer cells in the ascites of tumor-bearing mice, we next sought to determine the mechanism by which β-glucan eliminates cancer cells from the peritoneal fluid.

One distinguishing characteristic of peritoneal resident macrophages (PRMΦs) is their ability to rapidly aggregate around foreign particles or pathogens, entrapping them in clot-like structures and facilitating their clearance from the peritoneal fluid (Barth et al., 1995). This process is known as the MΦ disappearance reaction (MDR) and is critical to control infection in the peritoneal cavity (Vega-Pérez et al., 2021; Zhang et al., 2019). To test whether MDR could be responsible for cancer cell clearance, we administered GFP-labeled OvCa cells concurrently with β-glucan and analyzed peritoneal lavage 5 hours later (Figure S2A). We first confirmed MDR in our model by observing the disappearance of PRMΦs (F4/80^hi^CD11b^hi^ICAM2^hi^) in the peritoneal fluid following β-glucan administration (Figure 2A). We next looked for the presence of GFP^+^ cancer cells in the peritoneal fluid to see if they would also “disappear.” Indeed, the presence of both ID8 cells (Figure S2B) and KPCA cells (Figure 2B and S2C) were significantly reduced in β-glucan-treated mice. Because cancer cell clearance does not appear to be dependent on cell type, KPCA cells were used for the remaining experiments. Similarly to what has been previously reported in infection models, clot-like structures also formed in our cancer model following β-glucan treatment. These structures could be visualized as ∼1-2mm GFP^+^ clots freely floating in the peritoneal cavity (Figure 2C & S2D) and flow cytometric analysis of these structures confirmed the presence of CD45^-^GFP^+^ KPCA cells within (Figure 2D), thus suggesting that MDR can indeed capture cancer cells in the peritoneal fluid. To confirm this, we examined the peritoneal lavage of mice treated with clodronate-loaded liposomes (CLL) which deplete PRMΦs (van Rooijen and Hendrikx, 2010). Indeed, KPCA clearance was partially impaired in CLL-treated mice (Figure S2E). Moreover, concurrent administration of β-glucan with heparin, an anticoagulant which inhibits MDR by disrupting PRMΦ clotting (Zhang et al., 2019), also partially impaired KPCA clearance in the peritoneal lavage (Figure 2E). Additionally, OvCa cells captured within these clots were more apoptotic than untreated cancer cells freely floating in the peritoneal fluid as determined by TUNEL staining analyzed by flow cytometry (Figure S2F). Taken together, this data confirms that intraperitoneal administration of β-glucan can capture OvCa cells floating in the peritoneal fluid into clot-like structures by activating MDR, leading to elimination of OvCa cells from the peritoneal fluid.

**Figure 2.**
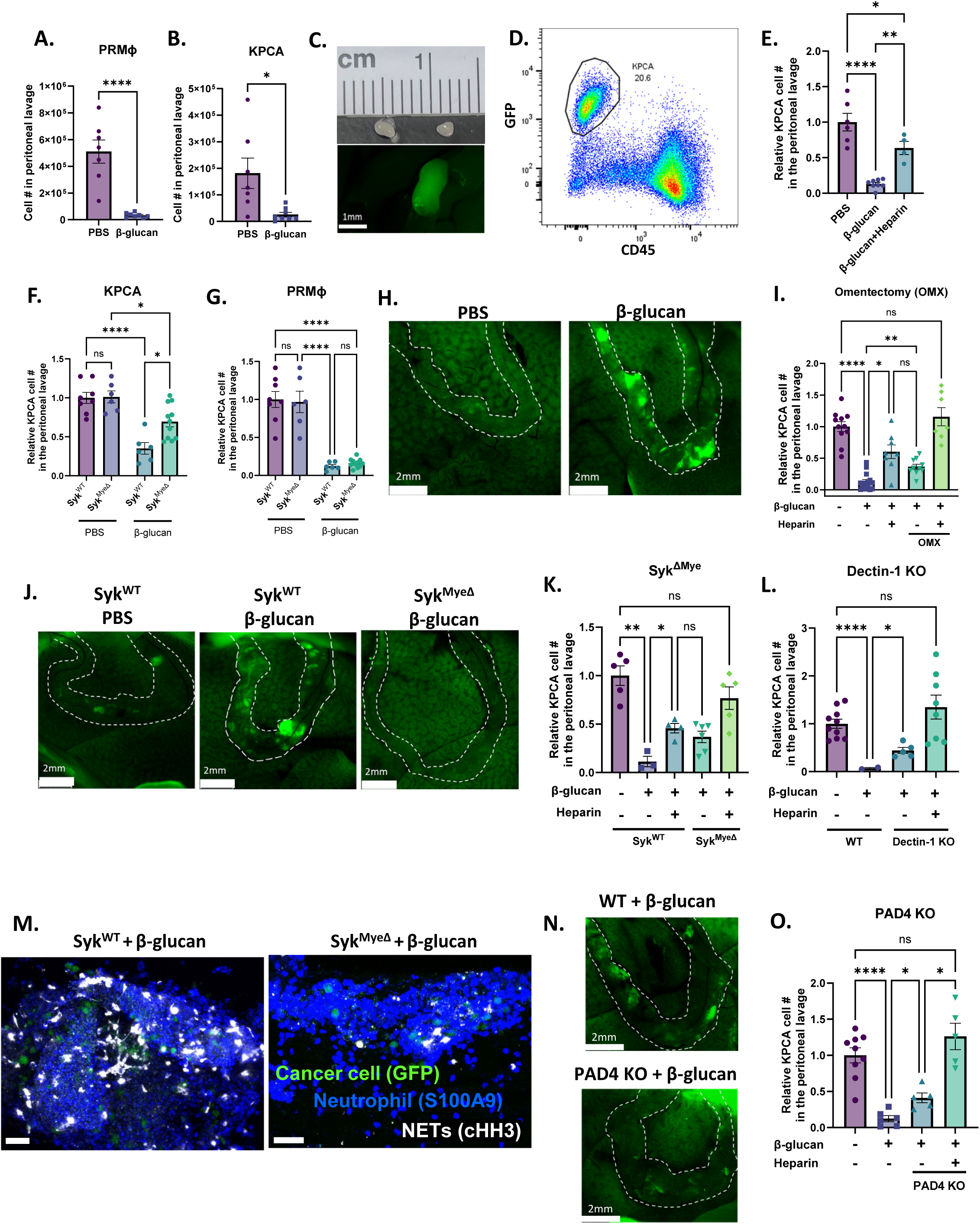
β-glucan captures OvCa cells into solid nodular structures via intraperitoneal clotting and Dectin-1-Syk-dependent NETosis in the omentum. Quantification of (A) peritoneal resident macrophages (PRMΦ) and (B) KPCA cells in the peritoneal lavage of mice 5 hours following PBS or β-glucan treatment. Representative (C) image and (D) flow plot of peritoneal clots formed in the peritoneal fluid following β-glucan treatment containing GFP+CD45-KPCA cells. (E) Quantification of PRMΦ and KPCA cells in the peritoneal lavage 5 hours after PBS, β-glucan, and β-glucan+heparin treatment. Quantification of (F) KPCA and (G) PRMΦ in the peritoneal lavage of Syk^WT^ and Syk^MyeΔ^ mice treated with PBS or β-glucan. (H) Representative images of omentum in mice 5 hours after PBS and β-glucan treatment. Omentum were stretched over the liver for better imaging. (I) Quantification of KPCA cells in the peritoneal lavage in intact and omentectomized (OMX) mice treated as indicated with PBS, β-glucan, and heparin. (J) Representative images of omentum in Syk^WT^ and Syk^MyeΔ^ mice 5 hours after β-glucan treatment. Quantification of KPCA in the peritoneal lavage of (K) Syk^MyeΔ^ and (L) Dectin-1 KO mice 5 hours following treatment as indicated with PBS, β-glucan, and heparin. (M) representative confocal images of omentum of Syk^WT^ and Syk^MyeΔ^ mice 5 hours after β-glucan treatment. Positive cells were stained blue (S100A9; neutrophil), green (GFP; cancer cell), and white (cHH3; NETs). (N) representative images of omentum and (O) quantification of KPCA cells in peritoneal lavage in WT and PAD 4KO mice 5 hours after indicated treatment. One-way ANOVA and student’s t test were used. *p<0.05; **p<0.01; ****p<0.0001. Error bars are standard errors of the mean. Relative cell number is reported as fold change to the average of the control and was used when experimental replicates were combined.

We next wanted to investigate the molecular mechanism underpinning β-glucan-mediated cancer clearance from the peritoneal fluid. Dectin-1, a C-type lectin receptor expressed by myeloid cells, recognizes β-glucan and signals via downstream spleen tyrosine kinase (Syk) (Brown et al., 2002). To determine the contribution of Dectin-1-Syk signaling in cancer cell clearance, we first analyzed the peritoneal lavage of mice with Syk-deficient myeloid cells (Syk^MyeΔ^). Indeed, cancer cell clearance from the fluid was impaired in Syk^MyeΔ^ mice as compared to their littermate controls (Syk^WT^, Figure 2F). Surprisingly, impaired OvCa cell clearance occurred independent of MDR as PRMΦs were still undetectable in the peritoneal lavage of Syk^MyeΔ^ mice after β-glucan treatment (Figure 2G). Moreover, neither Syk deficiency nor MDR inhibition completely reversed OvCa cell clearance from the peritoneal fluid (Figure 2E & F), thus implying the existence of two independent cancer clearance mechanisms: one which involves MDR and one which requires Syk signaling in myeloid cells.

To further elucidate the MDR-independent, Syk-dependent mechanism, we chose to focus on the omentum in β-glucan treated mice. Known as the “policeman of the abdomen,” the omentum is another key player critical for clearing peritoneal contaminants and coordinating protective immune responses during peritonitis (Català et al., 2022; Meza-Perez and Randall, 2017). Notably, zymosan (a type of β-glucan) has been reported to be rapidly sequestered in the omentum following i.p. injection (Jackson-Jones et al., 2020). To test whether the omentum can also capture OvCa cells following intraperitoneal β-glucan administration, we imaged the omentum *in situ* 5 hours after injecting KCPA cells and β-glucan. Mouse body cavities were opened, and the omentum was gently stretched across the liver to ensure low background fluorescence and clear imaging. Green fluorescing OvCa cells were clearly visualized in the omentum following β-glucan treatment (Figure 2H), indicating that the omentum can indeed sequester OvCa cells following β-glucan administration. Moreover, OvCa cell clearance from the peritoneal lavage was partially but significantly reversed in mice whose omentum were surgically removed by omentectomy (OMX, Figure 2I). Similar to what was observed in Syk^MyeΔ^ mice, reversal of OvCa cell clearance occurred in OMX mice independent of MDR (Figure S2G), thus supporting the notion that clearance by the omentum and MDR likely occurs independent of one another. To confirm this, we administered heparin in OMX mice where both MDR and omentum capture was inhibited and observed cancer cell clearance from the peritoneal fluid was completely reversed in these mice (Figure 2I). This supports the existence of two complementary pathways in two independent structures (peritoneal fluid and the omentum) that are required for total OvCa cell clearance by β-glucan.

Given that cancer clearance in Syk^MyeΔ^ mice phenocopied what was observed in OMX mice, we next wanted to test whether Dectin-1-Syk signaling could drive cancer cell sequestration in the omentum. Indeed, fewer cancer cells were observed in the omentum of Syk^MyeΔ^ mice as seen by imaging and flow cytometry (Figure 2J & S2H). Moreover, like OMX mice, heparin administration in Syk^MyeΔ^ and constitutive Dectin-1 knockout mice (Dectin-1 KO) once again completely reversed OvCa cell clearance in the peritoneal lavage (Figure 2K & L). Moreover, depletion of MΦs using CLL in Syk^MyeΔ^ mice also completely reversed OvCa cell clearance (Figure S2I), further confirming that Dectin-1-Syk signaling drives OvCa cell capture in the omentum independent of MDR.

Finally, we sought to identify the mechanism through which OvCa cells were trapped within the omentum. A recent report demonstrated that the capture of zymosan in the omentum is mediated by neutrophil recruitment and activation of neutrophil extracellular traps (NETs) (Jackson-Jones et al., 2020). Additionally, Syk signaling has been reported to be a master regulator of NETosis in response to β-glucan (Nanì et al., 2015; Negoro et al., 2020; Zhu et al., 2023). Given that Syk signaling in myeloid cells is crucial for OvCa cell capture in the omentum, we posited that OvCa cells could be captured via Syk-dependent NETosis in the omentum. Whole-mount confocal imaging of the omentum 5 hours after β-glucan and KPCA administration demonstrated that NETs, marked by citrullinated histone H3 (cHH3), were indeed activated by β-glucan and localized with neutrophils and OvCa cells in the omentum (Figure 2M). Moreover, this signal decreased in Syk^MyeΔ^ mice (Fig. 2M), confirming that Syk-mediated NETosis may drive OvCa cell capture by the omentum. Finally, we found that inhibiting NETs directly by genetically deleting peptidyl arginine deiminase type IV in mice (PAD4 KO) (Li et al., 2010) partially reversed OvCa cell capture by the omentum (Figure 2N) and that adding heparin in PAD4 KO mice once again fully reversed OvCa cell clearance in the peritoneal lavage (Fig. 2O). Thus, β-glucan induces capture of OvCa cells by the omentum via Dectin-1-Syk-mediated NETosis.

Taken together, we identified two nonredundant yet complementary pathways that mediate clearance of disseminating OvCa cells from the peritoneal fluid by β-glucan: (1) an MDR-mediated intraperitoneal clotting mechanism in the peritoneal fluid and (2) a Dectin-1-Syk-dependent NETosis mechanism in the omentum. A simplified illustration of these pathways can be found in Figure S2J.

### Combining β-glucan with IFNγ reduces KPCA tumor burden through host immunity

Although β-glucan treatment alone effectively controlled fluid tumor burden, it was unable to control metastatic growth of KPCA tumors *in vivo* (Figure 1H). Emerging evidence suggests that therapies combining IFNγ and pathogen-derived molecules can be utilized against cancers and have been tested in a few clinically relevant, orthotopic cancer models (Sun et al., 2021; Wattenberg et al., 2023). In light of this, we adopted a similar approach and posited that β-glucan in combination with IFNγ could potentially treat KPCA tumors.

To test the efficacy of β-glucan+IFNγ (BI) against KPCA tumors, 1×10^6^ luciferase-tagged KPCA cells were seeded i.p. and treated as shown in Figure 3A. BI significantly reduced KPCA tumor burden as visualized by bioluminescence imaging, flow cytometry, and omentum weight (Figure 3B, S3A & S3B), but single agent β-glucan or IFNγ did not. Reduction of KPCA tumor burden was completely impaired in IFNγ receptor knockout mice (IFNγR KO, Figure 3C) and BI treatment did not reduce KPCA number *in vitro* (Figure S3C), indicating that the therapy is not directly cytotoxic to tumors but requires cancer-extrinsic IFNγR-mediated host immune responses. MΦ activation can promote tumor killing by activating T cells (Sun et al., 2021). To test whether T cells are required for the antitumor activity of BI, we depleted T cells *in vivo* using monoclonal antibodies against CD4 and CD8 (αCD4+αCD8). Indeed, mice lacking CD4^+^ and CD8^+^ T cells are unable to control tumor burden following BI (Figure 3D). Taken together, this data indicates that BI activates host immunity to robustly control metastatic KPCA tumors *in vivo*.

**Figure 3.**
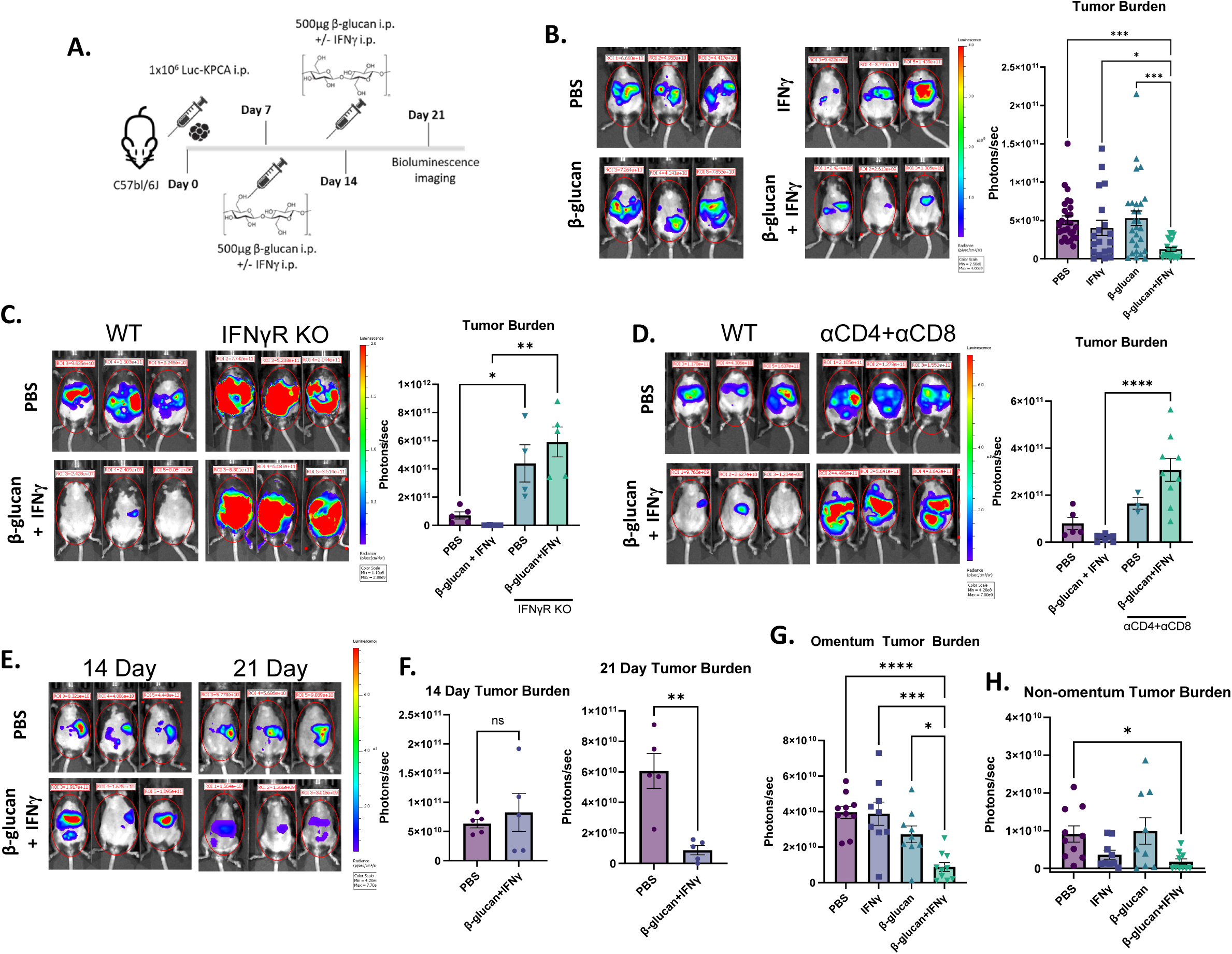
Combining β-glucan with IFNγ reduces KPCA tumor burden through host immunity. (A) β-glucan+IFNγ (BI) treatment timeline in the KPCA OvCa model. (B) Representative bioluminescence images and quantification of bioluminescence signals in mice treated with PBS, IFNγ, β-glucan, or BI. Representative bioluminescence images and quantification of bioluminescence signals in (C) IFNγ Receptor knockout mice and (D) T cell-depleted mice treated with PBS or BI. (E) Representative bioluminescence images and (F) quantification of bioluminescence signals of PBS and BI treated mice 14- and 21-days post tumor seeding. Quantification of (G) Omentum and (H) non-omentum body cavity bioluminescence signals in PBS-, IFNγ-, β-glucan-, or BI-treated mice. One-way ANOVA and student’s t test were used. *p<0.05; **p<0.01; ****p<0.0001. Error bars are standard errors of the mean.

We next sought to determine whether BI could drive tumor regression. To this end, we tracked tumor growth longitudinally following treatment via bioluminescence imaging (Figure S3D). Notably, there was no difference in tumor burden across all treatment groups on days 8, 10, or 14, however there was a significant decrease in tumor burden in only BI-treated mice on day 21 (Figure 3E, 3F, and S3E). These data indicate that BI may induce direct tumor killing and disease regression, although it does not rule out the possibility that BI may reduce tumor growth. Compartmental imaging of the omentum and the non-omentum body cavity (Figure S1D) revealed that the omentum had the greatest tumor burden and that both β-glucan and IFNγ were required to control tumors in the omentum and throughout the rest of the cavity (Figure 3G and 3H) Ascites accumulation was eradicated in both β-glucan- and BI-treated mice (S3F and S3G). To test the toxicity of BI we analyzed multiple organ damage markers in the serum of treated mice on day 21 and noticed no significant difference between PBS- and BI-treated mice, indicating its safety (Figure S3H). Moreover, there was no difference in body weight between PBS- and BI-treated mice (Figure S3I), however, more comprehensive toxicity analysis is required before testing in humans. Taken together, this data demonstrates that BI activates host immunity to robustly kill KPCA tumors in metastatic sites in the peritoneal cavity *in vivo*.

### BI enriches IL27^+^ antitumor MΦs in omentum tumors

BI can signal through receptors expressed by MΦs (Dectin-1 and IFNγR respectively). To test whether MΦs are required for BI-induced antitumor immunity, we depleted MΦs using clodronate-loaded liposomes (CLL) during the course of BI treatment and found that mice treated with CLL had a higher tumor burden than BI+PBS-treated mice (Figure 4A), suggesting MΦs are required for the optimal tumor control induced by BI.

**Figure 4.**
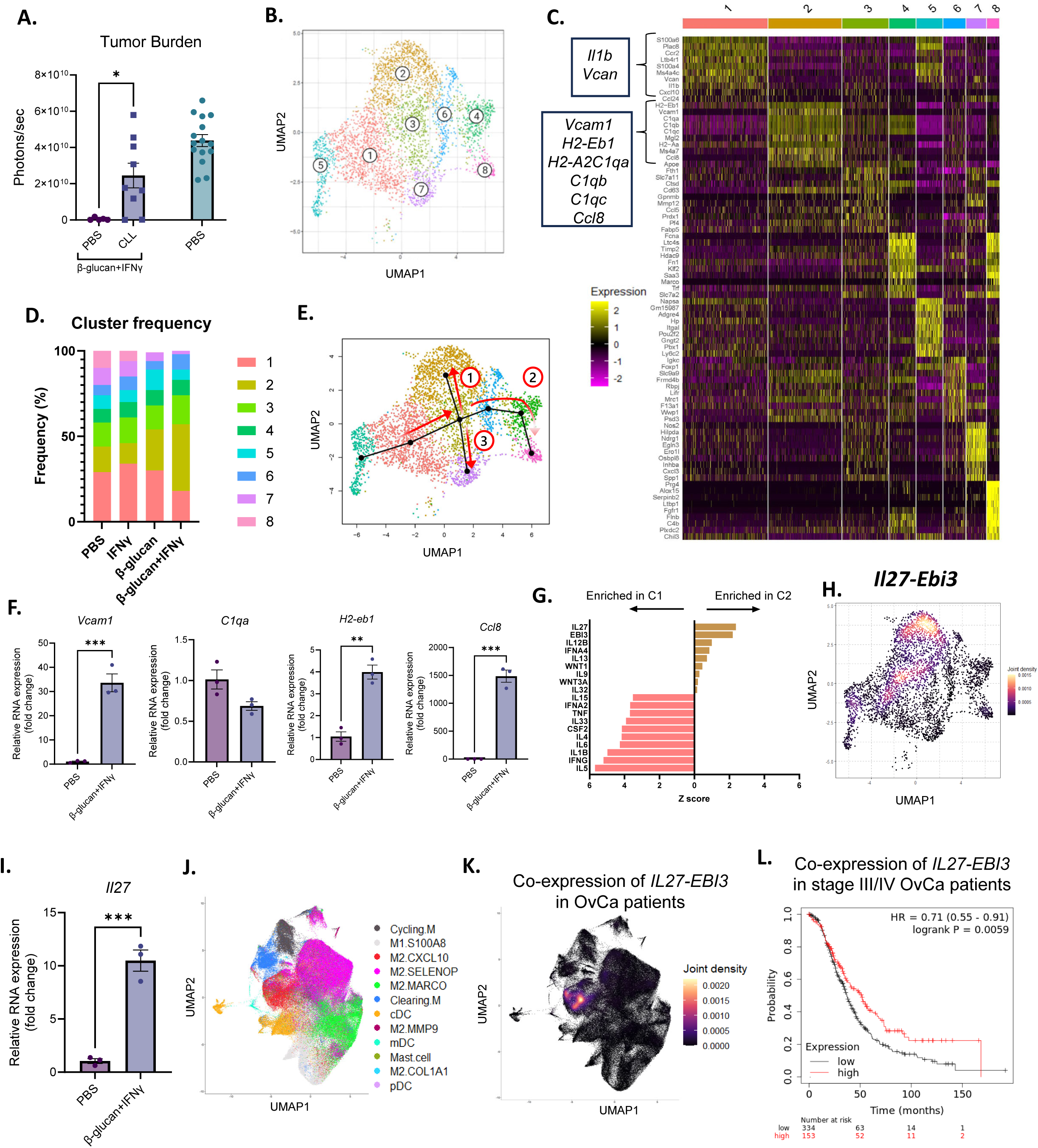
BI enriched IL27+ macrophages in omentum tumors. (A) Whole body bioluminescence signals of PBS or CLL-treated mice treated with BI and control mice. (B) A UMAP plot of monocyte/MΦ clusters in omentum tumors. (C) Top expressed genes in all monocyte/MΦ clusters. (D) Frequencies of eight identified monocyte/MΦ clusters in omentum tumors. (E) Slingshot trajectory analysis from the origin (Cluster 5) through three independent pathways (red arrows). (F) qPCR analysis of Cluster 2-specific genes in monocyte-derived MΦs treated with PBS or BI. (G) Top upregulated IPA cytokine regulators in Clusters 1 and 2. (H) *Il27* and *Ebi3* co-expression heatmap in monocyte/MΦ clusters. (I) *Il27* expression in monocyte-derived MΦs treated with PBS or BI analyzed by qPCR. (J) UMAP monocyte/MΦ clusters and (K) *IL27* and *EBI3* co-expression in tumors from human OvCa patients. (L) Overall survival analysis in late stage OvCa patients (stage III and IV) with high and low co-expression of *IL27-EBI3.* Student’s t test and log rank test were used. *p<0.05; **p<0.01; ****p<0.0001. Error bars are standard errors of the mean.

Next, we sought to investigate how MΦs facilitated antitumor immunity. We focused on omentum tumors because they were the largest tumor burden and very few tumor nodules were present in any other peritoneal compartment or the fluid following BI (Figure 3G and 3H). Flow cytometric analysis of omentum tumors revealed a reduction in pro-tumor Arginase-1^+^ and Tim-4^+^ MΦs (Figure S4A), which have been reported to promote tumor progression by suppressing T cell functions (Noy and Pollard, 2014; Xia et al., 2020). To determine whether there is a unique subset of monocytes/MΦs which may promote tumor antitumor activity, we performed scRNA-seq on omentum tumors and identified 8 clusters of monocytes/MΦs (Figure 4B-C). Using the Immunological Genome Project dataset as a reference, Clusters 1 and 5 were identified at monocytes; all other clusters were identified as MΦs. Notably, Cluster 2 frequency was selectively enriched in BI-treated mice, while Cluster 1 was reduced (Figure 4D). Cluster 1 monocytes selectively expressed inflammation-related genes such as *Il1b* and *Vcan*, whereas Cluster 2 MΦs were enriched with genes related to antigen presentation (*H2-Aa, H2-Eb1*), immune activation (*Ccl8*, *Vcam1*), and complement activation (*C1qa, C1qb, C1qc*) (Figure 4C).

To better understand the origin and development of Cluster 2 MΦs we performed Slingshot trajectory analysis of our scRNA-seq dataset. Setting Cluster 5 monocytes as the origin, we found three distinct differentiation pathways (Figure 4E). All trajectories passed from Cluster 5 through Clusters 1 and 3 before diverging and terminating in Clusters 2, 7 and 8. These data suggest that Cluster 2 MΦs developed from monocytes through a unique pathway. To support this hypothesis, we cultured monocytes isolated from bone marrow and stimulated them with BI during their *in vitro* differentiation into MΦs. MΦs which differentiated in the presence of BI upregulated multiple markers of Cluster 2 MΦs, such as *Vcam1*, *H2-eb1*, and *Ccl8,* but not *C1qa* (Figure 4F). Thereby supporting the notion that Cluster 2 MΦs may arise from BI-treated monocytes, but their full maturation may require additional signals from the tumor microenvironment. Additionally, we investigated whether BI could change bone marrow progenitors and monocytes *in vivo*. 1 week after BI injection, we analyzed bone marrow cells by flow cytometry and found that Lin-Sca1+cKit+ progenitor cells (LSK), long-term hematopoietic stem cells (LT-HSC), multipotent progenitor cells (MPP), and Ly6C^hi^ monocytes were all increased compared to PBS-treated mice (Figure S4B). This is consistent with recently reported β-glucan-induced immune training (Ding et al., 2023; Kalafati et al., 2020; Mitroulis et al., 2018). These data together suggest that Cluster 2 MΦs develop from bone marrow monocytes.

To understand what signals Cluster 2 MΦs may upregulate to activate T cells to drive an antitumor response, we chose to focus on MΦ-derived cytokines, which are key approaches MΦs use to activate T cells. Ingenuity Pathway Analysis (IPA) specifically focusing on cytokine regulators in Cluster 1 and 2 confirmed the pro-inflammatory phenotype of Cluster 1 monocytes as inflammatory cytokine pathways were highly enriched, such as IL1β and IL6 (Figure 4G). In contrast, both subunits of the IL27 cytokine heterodimer (*Il27* and *Ebi3*) were significantly enriched in Cluster 2 (Figure 4G), and *Il27* and *Ebi3* co-expression was predominately detected in Cluster 2 MΦs (Figure 4H). Moreover, monocyte-derived MΦs in the presence of BI expressed higher levels of *Il27* when compared to untreated MΦs (Figure 4I). IL27 is an unconventional IL12-family cytokine that has both pro- and antitumor properties depending on tumor types (Fabbi et al., 2017; Yoshida and Hunter, 2015), while IL12 is the classical MΦ-derived antitumor cytokine (Chan et al., 1991; Kaczanowska et al., 2021). Surprisingly, IL12 was almost undetectable in any of our MΦ subpopulations (Figure S4C).

Lastly, to determine whether IL27 signaling is relevant in OvCa patients, we analyzed a recently published scRNA-seq dataset of tumors and ascites from patients with metastatic OvCa (Vázquez-García et al., 2022). In these tumors, *IL27* and *EBI3* were exclusively co-expressed in only monocytes/MΦs (Figure S4D & S4E), specifically in the ‘M2.CXCL10’ subset (Figure 4J & 4K). Interestingly, these ‘M2.CXCL10’ MΦs are characterized by high expression of *CCL8*, which is one of the most differentially expressed genes in Cluster 2 MΦs identified in our dataset (Figure 4C), indicating that transcriptional regulators that control IL27^+^ MΦs are conserved between mice and humans. To our surprise, we did not find significant expression of IL12 in these tumor samples (Figure S4F), consistent with the result from our mouse model. Finally, utilizing public datasets, we analyzed the correlation between overall patient survival and IL27 expression. Indeed, higher co-expression of *IL27* and *EBI3* significantly correlated with improved survival in patients with stage III/IV metastatic OvCa (Figure 4L, p=0.0059). In summary, these results suggest that BI treatment causes regression of metastatic OvCa by promoting differentiation of monocytes into IL27^+^ antitumor MΦs.

### IL27 contributes to BI treatment by activating T cells and is specifically secreted by BI-stimulated MΦs

To test whether IL27 contributes directly to the antitumor response of BI, we neutralized IL27 using a monoclonal antibody against IL27p28 in the presence of BI *in vivo*. Indeed, IL27 neutralization significantly impaired the antitumor activity in both omental and mesenteric metastases as compared to IgG controls (Figure 5A & 5B). Given the requirement of T cells for BI efficacy (Figure 3D), we next evaluated changes in T cells following BI treatment and examined the potential role of IL27 underlying these responses. T cells from PBS- and BI-treated omentum tumors were restimulated *ex vivo* and analyzed by flow cytometry. T cells secrete cytokines such as IFNγ and TNF upon activation to drive anti-tumor immune responses. In BI-treated tumors, although CD8^+^ T cell frequencies did not change (Figure S5A), IFNγ^+^ and TNF^+^ CD8^+^ T cells were enriched (Figure S5B) and IFNγ and TNF mean fluorescent intensity (MFI) was increased (Figure 5C). Interestingly, in CD4^+^ T cells, only the frequency of TNF^+^ was increased, and no other significant changes were observed (Figure S5C). Therefore, BI treatment is immunostimulatory and its efficacy likely relies on the activation of CD8^+^ cytotoxic T cells.

**Figure 5.**
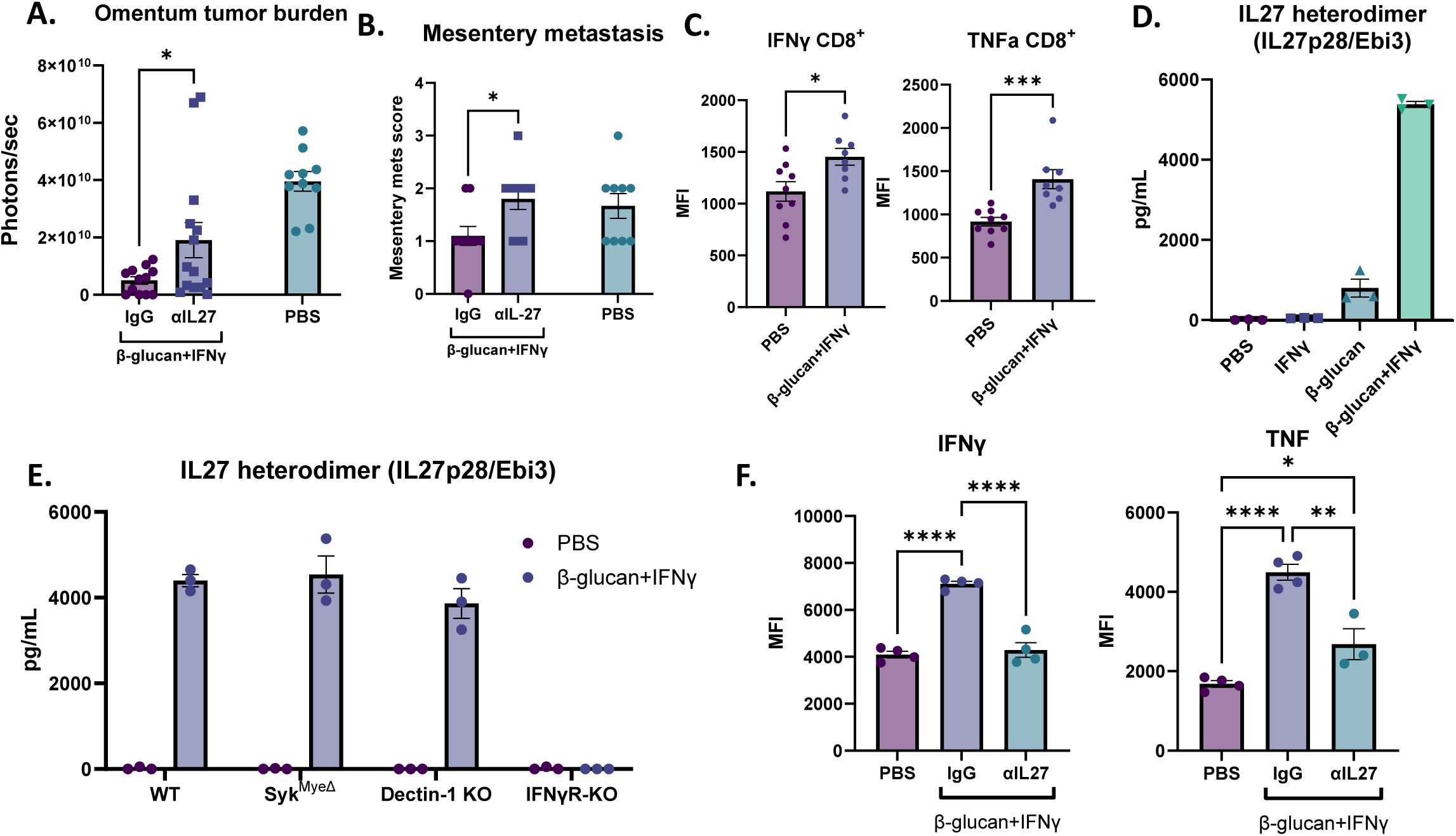
IL27 contributes to BI treatment by activating T cells and is specifically secreted by BI-stimulated MΦs. (A) Bioluminescence signals of omentum tumors and (B) mesentery metastasis scores from mice injected with IgG or αIL27 treated with BI as well as PBS-treated control mice. (C) Mean fluorescent intensity (MFI) of IFNγ and TNF in CD8^+^ T cells in omentum tumors from control or BI treated mice analyzed by flow cytometry. (D) ELISA quantification of IL27 heterodimer in supernatant from BMDM stimulated with PBS, IFNγ, β-glucan, or BI. (E) ELISA quantification of IL27 heterodimer in supernatant from WT-, Syk^MyeΔ^, Dectin-1 KO, and IFNγR KO BMDM cultured with PBS or BI. (F) IFNγ and TNF MFI of CD8^+^ T cells cocultured with MΦ pretreated with PBS or BI in the presence of αIL27 antibody or control IgG. Student’s t test and One-way ANOVA were used. *p<0.05; **p<0.01; ***p<0.001; ****p<0.0001. Error bars are standard errors of the mean.

To test whether MΦ-derived IL27 contributes to the immunostimulatory effect seen in BI treatment, we established an *in vitro* system using MΦs and CD8^+^ T cells. First, to test whether the IL27 can be directly secreted from MΦs stimulated by BI, we treated bone marrow derived MΦs (BMDMs) with BI for 48 hours and tested the supernatant for secreted IL27 using an IL27/EBI3 heterodimer-specific ELISA. Interestingly, while β-glucan alone can generate a small amount of IL27, both β-glucan and IFNγ are required for robust IL27 secretion (Figure 5D). It has been previously reported that the p28 subunit of IL27 (also known as IL30 when IL27p28 forms a monomer) can be generated in MΦs stimulated with IFNγ or LPS (another PAMP molecule) as detected by an IL27p28 ELISA (not specific for IL27 heterodimer). In this context, a single agent is sufficient to induce IL27p28 production, but combining the two agents synergizes to produce maximal amount of IL27p28 (Liu et al., 2007). Notably, we see a similar pattern in IL27p28 secretion in BMDMs treated with β-glucan or IFNγ (Figure S5D). Therefore, secretion of the IL27 heterodimer is specific to BI treatment despite IL27p28 generation induced by single agents, suggesting that both agents may be necessary for the transcription of EBI3, but how this occurs remains unknown. Surprisingly, IL27 did not require Dectin-1-Syk signaling as BMDMs from Dectin-1 KO and Syk^MyeΔ^ mice still secreted IL27 following BI treatment (Figure 5E). On the other hand, the receptor for IFNγ was crucial for IL27 generation as loss of this receptor completely ablated IL27 secretion in BMDMs (Figure 5E). Therefore, we show that secretion of the IL27 heterodimer is a specifically regulated event which requires IFNγ receptor signaling but not Dectin-1 or Syk. Given the role IL27 plays in BI efficacy (Figure 5A), the specific generation of IL27 by BI may offer one explanation as to why single agent β-glucan or IFNγ treatment was not sufficient in controlling OvCa tumor burden (Figure 3B).

To test whether MΦ-derived IL27 could drive the immunostimulatory phenotype in CD8^+^ T cells, naïve OT-I CD8^+^ T cells were co-cultured with PBS- or BI-pretreated BMDMs, OVA peptide, and dendritic cells in the presence of control or an IL27p28 neutralization antibody. Indeed, co-culturing T cells with BI-pretreated BMDMs increased the frequency of IFNγ^+^, TNF^+^, and IFNγ^+^TNF^+^ CD8^+^ T cells (Figure S5E and S5F) and increased IFNγ and TNF mean fluorescence intensity (MFI) (Figure 5F) in an IL27-dependent manner. BI-pretreated MΦs also increased Granzyme B expression (a cytotoxic effector molecule in CD8^+^ T cells) but did so independent of IL27 (Figure S5G). Taken together, these data suggest that BI treatment directly stimulates IL27 expression in MΦs and that MΦ-derived IL27 promotes the anti-tumor activity of BI treatment via activating cytotoxic CD8^+^ T cells.

### BI extends overall survival in both chemoresistant and chemo-sensitive models and dramatically enhances chemotherapy response in the chemo-sensitive model

To test whether β-glucan+IFNγ can be combined with a platinum-based chemotherapy (standard-of-care for OvCa patients) to treat metastatic OvCa, we treated the homologous recombination (HR)-proficient chemoresistant KPCA and HR-deficient chemo-sensitive BPPNM tumors (Iyer et al., 2021) with BI once a week for two weeks with or without carboplatin and monitored their overall survival. As expected in the chemoresistant KPCA model, carboplatin alone did not yield any survival advantage compared to PBS controls, but BI significantly extended the overall survival and led to 20% cure. In addition, combining BI with carboplatin did not lead to a survival advantage compared to BI alone. On the other hand, in the HR-deficient chemo-sensitive BPPNM model, carboplatin alone significantly extended the overall survival, while BI modestly prolonged the survival (Figure 6B). In sharp contrast to the KPCA model, combining BI with carboplatin led to 80% cure of BPPNM tumors (Figure 6B). These results suggest that combining BI with platinum-based chemotherapy may yield a significant therapeutic advantage compared to chemotherapy alone in HR-deficient OvCa, while BI may be a treatment option for chemoresistant HR-proficient OvCa.

**Figure 6.**
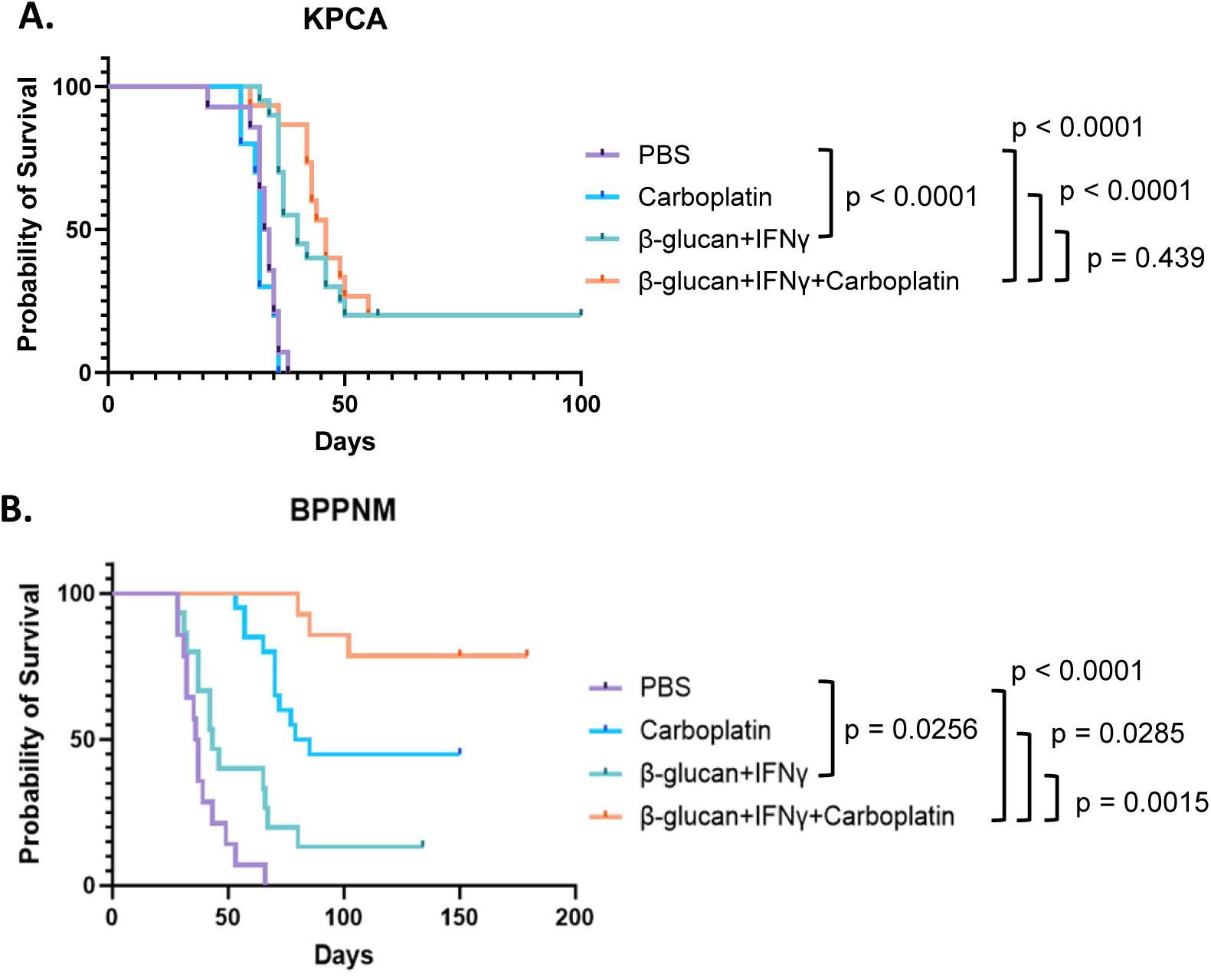
BI extends overall survival in both chemoresistant and chemo-sensitive models and dramatically enhances chemotherapy response in the chemo-sensitive model. (A) Survival curves of KPCA tumor-bearing mice treated with PBS, carboplatin, BI, or BI+carboplatin as indicated. The numbers of mice are PBS n=14, carboplatin n=10, BI n=20, BI+carboplatin n=15. The graph is a combination of three independent experiments. (B) survival curves of BPPNM tumor-bearing mice treated with BI and carboplatin as indicated. The numbers of mice are PBS n=14, carboplatin n=20, BI n=15, BI+carboplatin n=14. The graph is a combination of three independent experiments. Log-rank test was used.

## Discussion

Despite incredible advancements in the treatment of other cancers due to the rise of T cell-based immunotherapies, treatments for metastatic OvCa have not improved. In this study, we present an alternative immune therapy which harnesses myeloid cells against aggressive metastatic OvCa. In summary, we report here that β-glucan, a pathogen associated molecular pattern (PAMP) molecule, in combination with the immunogenic cytokine IFNγ coordinates a robust antitumor response in a patient-relevant murine model of metastatic OvCa. Tumor burden regressed in multiple metastatic compartments including the peritoneal fluid, omentum, and other parts of the peritoneal cavity. Surprisingly, β-glucan alone was sufficient to control fluid tumor burden despite not controlling solid tumor progression. Mechanistically, we demonstrated that sequestration of tumor cells out of the peritoneal fluid requires intraperitoneal clotting accompanied with MDR and Dectin-1-Syk-dependent NETosis in the omentum (Figure S2J). Systemic tumor regression following BI was facilitated through tumor-extrinsic signaling, requiring both T cells and IFNγR signaling from the host. ScRNA-seq analysis of omentum tumors revealed a selective enrichment of monocyte-derived IL27^+^ MΦs following BI. Neutralizing IL27 *in vivo* significantly impaired BI-induced antitumor immunity, demonstrating that IL27 indeed drives antitumor immunity likely through MΦs. In agreement with this notion, MΦs treated directly with BI *in vitro* can secrete IL27 and activate CD8^+^ T cells in an IL27-depdendent manner. Moreover, in patients with metastatic OvCa, *IL27/EBI3* co-expression predicted better overall patient survival. Finally, BI significantly extended overall survival of mice with clinically relevant metastatic OvCa and dramatically enhanced the efficacy of platinum-based chemotherapy *in vivo* in a clinically relevant, HR-deficient metastatic OvCa model. Taken together, these data suggest the therapeutic potential of BI in treating metastatic OvCa in patients.

In contrast to the hematological route of metastasis seen in other cancers, OvCa cells readily disseminate directly into the peritoneal fluid as single cells or multicellular aggregates prior to seeding in secondary peritoneal metastatic sites. This route of dissemination is uniquely challenging as these fluid-bound spheroids are functionally dormant, rendering them resistant to anoikis (a form of apoptosis following cell detachment) and proliferation-targeting chemotherapy (Shepherd and Dick, 2022). At the same time, highly invasive and therapy resistant OvCa stem cells have been reported to be enriched in malignant ascites in patients (Raghavan et al., 2019; Ward Rashidi et al., 2019). One theory of relapse suggests that cancer cells left behind in the fluid following surgical debulking and therapy give rise to chemoresistant patient relapse (Liao et al., 2014; Shepherd and Dick, 2022). Therefore, numerous studies have attempted to elucidate cancer intrinsic vulnerabilities of disseminating cancer cells in the peritoneal fluid in an effort to improve therapy response (Buensuceso et al., 2020; Haagsma et al., 2023; Latifi et al., 2012). To our knowledge, this study is the first of its kind to demonstrate a cancer *extrinsic* approach for targeting these cells. Here we demonstrate that peritoneal immune responses which canonically target pathogens can be exploited to target disseminating cancer cells in the peritoneal fluid, seemingly independently of cell types (Figure 2B and S2B). Moreover, not only did intraperitoneal β-glucan administration sequester cancer cells out of the peritoneal fluid, it also induced acute cancer killing in peritoneal clots (Figure S2F). Interestingly, BI enhanced cancer killing in these clots as well (Figure S5H and S5I). The consequence of clot biology in killing cancer cells remains an open question and warrants further study. Meanwhile, although the acute fate of cancer cells trapped by the omentum was not investigated in this study, the consequence of rapid sequestration of cancer cells into the omentum still has the potential to revolutionize OvCa treatment. Despite efforts in improving therapeutics against OvCa, one of the greatest predictors of prognosis in stage III and IV patients is still the maximal removal of tumors during cytoreduction surgery (Bristow et al., 2023). Because most OvCa patients will undergo omentectomy as part of treatment, preoperative intraperitoneal administration of β-glucan could potentially improve surgical outcomes by trapping fluid cancer cells in the organ prior to its removal. However, further research is needed to test the efficacy and safety of such an approach. Still, this study shows for the first time that exploring cancer extrinsic mechanisms of cancer targeting in the peritoneal fluid holds promise to address a critical need.

The pleiotropic cytokine IL27 has been reported to contribute to tumor immunity in a context-dependent manner (Fabbi et al., 2017). It was initially reported as an IL12-like cytokine secreted by antigen-presenting cells that drives T cell activation (Pflanz et al., 2002) and antitumor immunity (Liu et al., 2022; Patidar et al., 2022). More recently, accumulating evidence demonstrates the role of IL27 in promoting expansion of regulatory T cells (Do et al., 2017; Hall et al., 2012), expression of T cell checkpoint receptors (Carbotti et al., 2015; Hirahara et al., 2012), and survival of tumor cells (Jia et al., 2016), revealing its context-dependent role in modulating tumor immunity. This starkly contrasts to IL12, which is universally considered to be antitumor. IL12 is strongly induced by PAMP/IFNγ stimulation *in vitro* and is a hallmark of antitumor MΦs. Recent *in vivo* work has demonstrated that co-stimulation of MΦs by MPLA (a PAMP molecule, acting as a TLR4 agonist) and IFNγ in breast cancer and OvCa induces IL12 production by MΦs and promotes antitumor T cell activity (Sun et al., 2021). However, the serious toxicity of IL12 in humans prevented its clinical usage (Leonard et al., 1997). To this end, IL27 could be a promising cytokine that stimulates local rather than systemic antitumor immunity. Interestingly, in published human and our own mouse scRNA-seq datasets of OvCa, expression of IL12 was not highly induced (Figure S4C & S4F), suggesting that different tumor microenvironments can induce differential antitumor cytokine-secretion in MΦs. These suggest that IL27 is a promising antitumor cytokine in treating metastatic OvCa. Whether IL27 is a viable therapeutic target in OvCa requires future investigations. Finally, as to the regulation of IL27 expression and secretion, we found that both agents in BI treatment were required to stimulate IL27 heterodimer secretion *in vitro.* Meanwhile, either β-glucan or IFNγ on their own are sufficient to drive the secretion of IL27p28 (the monomer form is also known as IL30), although BI is still necessary to produce the highest response (Figure S5D). This is consistent with previous reports which demonstrated that LPS (another PAMP molecule) induces IL27p28 expression and secretion, and this effect was enhanced by the addition of IFNγ (Liu et al., 2007). Therefore, understanding the genetic regulation of EBI3, the other subunit of IL27, or the secretion of the IL27 heterodimer following BI treatment may ultimately dictate IL27 regulation in MΦs. Moreover, future studies will answer how IL27 is induced *in vivo* and definitively demonstrate its cellular source and target.

To our surprise, we identified two β-glucan-driven mechanisms which did not require canonical Dectin-1-Syk signaling; (1) MDR and MDR-mediated cancer cell capture (Figure 2G, 2K, 2L) and (2) IL27 stimulation in MΦs (Figure 5E). This is surprising because Dectin-1 is considered to be the primary receptor for β-glucan signaling in MΦs (Brown et al., 2002; Goodridge et al., 2011). To our knowledge, while Syk-independent β-glucan signaling has been identified (Gringhuis et al., 2009; Herre et al., 2004), how β-glucan signals in MΦs independent of Dectin-1 is not well understood. β-glucan can bind to complement and therefore may signal through complement receptor 3 (CR3) in addition to Dectin-1. However, this interaction has been predominately studied in neutrophils and its contribution to phagocytosis of β-glucan particles in MΦs (like what is used for this study) is insignificant (Goodridge et al., 2009). Therefore, identifying the mechanism through which β-glucan drives MΦ clotting and IL27 is ongoing.

The ability of the innate immune system to generate memory responses to secondary infectious or inflammatory responses is called immune training (Netea et al., 2016). Immune training can be induced by β-glucan and was initially thought to be only involved in infection and inflammation (Netea et al., 2011; Quintin et al., 2012). Recently, it has been appreciated that β-glucan-mediated immune training could be one approach for the treatment of several cancers, inducing robust antitumor activity and protecting against relapse in preclinical models (Ding et al., 2023; Geller et al., 2022; Kalafati et al., 2020; Woeste et al., 2023). Here we observed an increase in bone marrow progenitors 1 week after BI treatment (Figure S4B), hinting at a potential role of immune training. However, unlike published immune training studies, we found that T cells were indispensable for BI-induced antitumor response against metastatic OvCa. Future studies are warranted to comprehensively evaluate the role of immune training in BI treatment.

In summary, we have established an alternative immune therapy which combines β-glucan and IFNγ to drive regression of metastatic OvCa *in vivo*. While more work is to be done on elucidating specific antitumor mechanisms in MΦs and other cell types following β-glucan+IFNγ and its impact on hematopoiesis and immune training, our data suggest (1) a novel concept of targeting tumor cells floating in the fluid by “relocating” them to adherence to an immune-rich environment for better killing and (2) a novel role of IL27 in promoting antitumor immunity in metastatic OvCa. We acknowledge that the antitumor role of IL27^+^ MΦs are only partially responsible for tumor elimination. Mechanisms mediated by other cell types will be investigated in future studies. Moreover, we propose to further investigate BI because of its tremendous therapeutic potential, which could improve the lives and survival of metastatic OvCa patients. Finally, while most immunotherapeutic approaches focus on targeting adaptive immunity, our data highlights the importance of harnessing innate immunity in developing robust anti-cancer immunotherapies.

### Experimental Model and Subject Details Cell lines

The original ID8 cell line, derived from spontaneous *in vitro* malignant transformation of C57BL6 mouse ovarian surface epithelial cells (Roby et al., 2000), was modified to express GFP and firefly luciferase. KPCA, and BPPNM cells, which were recently generated and characterized (Iyer et al., 2021), were generously gifted to us by Drs. Robert Weinberg and David Pepin at the Whitehead Institute and the Mass General Research Institute, respectively. KPCA cells were previously modified to express GFP and firefly luciferase. We modified BPPNM cells to express firefly luciferase. All cell lines were cultured in Dulbecco’s modified Eagle’s medium (DMEM, Corning, 10-017-CV) supplemented with 4% FBS (Gibco, 16140-071), 1% penicillin/streptomycin, 1X Insulin-Transferrin-Selenium (ITS, Gibco, cat# 41400-045), and 2ng/mL mouse epidermal growth factor (mEGF) at 37°C supplied with 5% CO_2_. Cells were passaged no more than 5 times prior to injection into mouse peritoneal cavities.

### Animal models

C57BL/6J (WT, #000664), B6.129S7-*Ifngr1^tm1Ag^t/J* (IFNγRKO, #003288), B6.129S6-Clec7atm1Gdb/J (Dectin-1 KO, # 012337) and B6.Cg-Padi4^tm1.1Kmow^/J (PAD4 KO, #030315) mice were purchased from Jackson Laboratories. *Lyz2^Cre/+^;Syk^fl/fl^* (Syk^MyeΔ^) and their Cre negative littermate controls (Syk^WT^) mice were generated by crossing *Lyz2^Cre^* mice (Jackson, #004781) with *Syk^fl/fl^* mice, which were generously gifted to us by Dr. John Lukens from the University of Virginia. All mice were housed in individual microisolator cages in a rack system with filtered air in Wistar’s mouse barrier facility and provided with shelter and enrichment to reduce stress. Reducing stress in mice is critical as stress has been reported to impede anti-cancer immunotherapy (Yang et al., 2019). Anecdotally, we observed similar effects in our model. Mice were kept on a 12hr light-dark cycle and had access to food and water *ad libitum*. All animal procedures were performed in accordance with the Wistar Institutional Animal Care and Use Committee under protocol 201536. Genotyping was performed utilizing Transnetyx automated genotyping services.

## Method Details

### Tumor implantation, treatment, and survival

Cells were harvested with trypsin-EDTA (corning), washed in PBS, and injected intraperitoneally (i.p.) into mice. 3×10^6^ ID8 cells were injected in 100μL PBS into 8-10wk old female WT mice and allowed to seed for two weeks prior to β-glucan treatment. Mice were treated with 500µg sonicated whole β-glucan particles (Invivogen, tlrl-wgp) in PBS i.p. or PBS vehicle control once every other week for two weeks for a total of two injections (Figure S1A). Two weeks after the final dose of β-glucan, tumor burden was assessed by IVIS Spectrum Imaging (PerkinElmer). For KPCA tumors, 1×10^6^ KCPA cells were injected i.p. into WT or IFNγRKO in 200μL of a 1:1 matrigel:PBS mix (Matrigel Matrix Basement Membrane, Corning, 354234). Tumors grew for 1 week prior to treatment. Mice were treated with 500µg β-glucan, 20ng recombinant mouse IFNγ (Peprotech, 315-05), β-glucan+IFNγ, or PBS vehicle control once a week for two weeks (Figure S1B). One week after the final treatment, mice were imaged by IVIS. Importantly, β-glucan was sonicated intermittently on high for 15min immediately prior to injection to ensure thorough disruption of β-glucan aggregates. Recombinant mouse IFNγ was gently reconstituted in molecular grade, sterile H_2_O and diluted in PBS to its working concentration. To preserve IFNγ activity, solutions were handled gently to reduce the presence of bubbles and never vortexed. For macrophage depletion studies, mice were treated with 100μL of clodronate-loaded liposomes (CLL, Liposoma C-005) one week prior to cancer capture studies. For tumor studies, 100μL of CLL was injected i.p. 5, 9, 14 and 19 days after tumor seeding. On day 14, CLL was administered 4 hours prior to WI treatment. For T cell depletion studies, αCD4 (Leinco Tech, C2838) and αCD8 (Leinco Tech, C2850) monoclonal antibodies were injected 150μg each i.p. in WT KPCA tumor-bearing mice 3 days following cancer seeding and then once a week for 2 weeks for a total of 3 injections. For the IL27 neutralization studies, 200µg αIL27p28 monoclonal antibody (InVivoMAb BE0326) (Marillier et al., 2014) was injected 2 days prior to β-glucan+IFNγ treatment, at the same time as treatment, and then two times a week for two weeks following treatment for a total of 6 injections. For treatment of experiments with carboplatin, 10-30 mg/kg once a week of carboplatin was administered with or without β-glucan/IFNγ, starting at day 7 for 2 weeks. The dose and timing of β-glucan and IFNγ were the same as mentioned above. On day 21, mice were imaged by IVIS and further monitored for survival analysis.

### Bioluminescent Imaging

*In vivo* and *ex vivo* bioluminescence imaging was performed on an IVIS 50 (PerkinElmer; Living Image 4.3.1), with exposures of 1 s to 1 min, binning 2–8, field of view 12.5 cm, f/stop 1, and open filter. For *in vivo* imaging, D-Luciferin (Gold Biotechnology, 150 mg/kg in PBS) was injected into the mice i.p. and imaged 10min later. Mice were maintained under general anesthesia by continuous inhalation of 2-1.5% isoflurane in 60% oxygen. For *ex vivo* imaging, mice were injected with D-Luciferin and euthanized after 5min. Mice peritoneal cavities were exposed, and the omentum was excised. Mouse carcasses (“non-omentum”) and omentum were placed in the machine and imaged as shown in Figure S2D. The total photon flux (photons/s) was measured from regions of interest using the Living Image 2.6 program.

### Mesentery Metastasis Scoring

Mesentery metastasis score was calculated based on the following criteria. Whole disseminated mesenteric tumors were counted and each mouse score was determined as following:

0: no tumor was detected,
1: number of tumor nodules is less than 10,
2: 2: number of nodules is 10-30,
3: 3: number of nodules is over 30.

### Acute cancer cell clearance by β-glucan

2×10^6^ cancer cells in 100µL of PBS were injected i.p. into 8-12wk male and female WT, Syk^MyeΔ^, Syk^WT^, Dectin-1 KO, or PAD4 KO mice immediately followed by 500µg sonicated β-glucan in 100µL PBS. 5hr later, mice were euthanized, peritoneal lavage was taken, and the omentum were imaged as described below (Figure S2A). Clots were harvested 24hr after β-glucan injection and digested prior to analysis by FACS as described below. Peritoneal lavage and clots were analyzed by FACS for the presence of GFP^+^CD45^-^ cancer cells and F4/80^hi^ICAM2^hi^CD11b^high^ PRMΦs. The Omentum was imaged to identify the presence of GFP+ cancer cells. To reduce background fluorescence, the omentum was gently stretched over the liver and imaged. For mice treated with heparin, 100units/mouse of heparin was given in the same syringe as β-glucan. In the MΦ depletion model, mice were treated with 100µL of CLL 1 week prior to cancer cell capture assays. Omentectomized mice were allowed to recover for 4 weeks prior to entering this study. Data from combined experiments are presented as fold change. Fold change was calculated as *n ÷ avg of the contol* where n=experimental value.

### Omentectomy

Extended-release buprenorphine (3.25mg/kg) was given subcutaneously preoperatively for pain management. Surgical removal of the omentum was accomplished under general anesthesia by continuous inhalation of 2–3% isoflurane in 60% oxygen using a veterinary vaporizer. Aseptic techniques were performed to maintain sterility in the surgical field. 6-8wk old male and female mice were used. Mice were placed on a heating pad in a supine position. A midline incision was made in the region of the stomach and the greater omentum was carefully exposed. The omentum is a thin, elongated adipose tissue that is located under the stomach and between the spleen and pancreas. The omentum was removed via electrocautery to avoid bleeding and the midline incision was closed with absorbable sutures in two layers (first the peritoneal wall was closed and then the skin). Mice were immediately placed in a clean heated cage and monitored until awake. A liquid recovery diet was provided, and mice were monitored daily for 7 days for signs of infection. Removal of both the entire greater and lesser omentum results in malperfusion of the stomach and spleen and thus was not feasible. Mice were allowed to recover for 4 weeks before any further experimental procedures were performed.

### Collection of peritoneal lavage and tissue dissociation

Peritoneal lavage was collected by flushing the peritoneal cavity with 6ml FACS buffer (DPBS with 2mM EDTA and 0.1% BSA). Mice were gently massaged to ensure optimal collection of peritoneal cells. Peritoneal clots were collected 24h after β-glucan injection and omentum tumors were collected at the conclusion of each study. Clots and tumors were digested in the same way using a cocktail of 1mg/ml collagenase IV and 100 µg/ml DNase I, in 1-4mL DMEM with 10% FBS depending on tissue size. The tissues were minced into small pieces prior to digestion at 37°C for 30min shaking at 800rpm. Samples were then passed through a 70µm cell strainer to collect single-cell suspension to be analyzed by flow cytometry. If necessary, single cell suspensions were treated with 1X red blood cell lysis buffer (BD Bioscience, 555899) on ice for 10min.

### Flow cytometry

Single-cell suspensions were collected as described above prior to staining with primary conjugated antibodies at their indicated dilutions (supplemental Table 1). Surface stain antibodies were incubated with cells on ice for 30min, washed with 1mL FACS buffer (DPBS with 2mM EDTA and 0.1% BSA). Intracellular staining was carried out using True-Nuclear™ Transcription Factor Buffer Set (Biolegend, 424401) according to manufacturer instructions. Briefly cells were fixed for 45min-1hr at room temperature in the dark. Notably, KPCA cells lose GFP signal following fixation. Following fixation, cells were washed 1x in permeabilization buffer and then incubated in permeabilization buffer with intracellular stain antibodies overnight at 4°C. Counting beads (Biolegend, 424902) were added to each sample to determine cell numbers. Samples were analyzed on a BD FACSymphony™ A3 Cell Analyzer using FlowJo software. For IFNγ and TNF staining in T cells, single cells from the omentum tumor were incubated for 4 hours with Cell Activation Cocktail (with Brefeldin A) (catalog no. 423303, Biolegend) prior to staining.

For bone marrow progenitors staining, 2-5 ×10^6^ Bone Marrow cells were stained as single cells suspension in FACS buffer for 100 min. at 4°C as follows. Cells were incubated with Aqua Live/Dead (BV510) and anti-CD16/32 BV711. After 10 min. the anti-CD34 PE was added. After 30 min., the remaining antibodies cocktail was added (anti-LY6G BUV563, B220 FITC, CD90.2 FITC, NK1.1 BUV661, Sca1 PerCpCy5.5, CD117 (cKit) BV786, CD135 BV4221, CD150 PECy5, CD48 AF700, LY6C BV605, CD81 PECy7, CD115 PE/Dazzle, CD11B BUV805, CD106 BUV737). Cells were washed and resuspended in 500 uL of FACS Buffer and acquired with a BD FACSymphony™ A5 Cell Analyzer and analyzed using FlowJo software.

### Fluorescence imaging

5 hours after the injection of β-glucan and KPCA cells, the omentum was excised and fixed in 4% paraformaldehyde overnight at 4 °C. On the next day, the tissues were rinsed with 3x PBS, blocked with 3% BSA and 1% Triton in PBS at room temperature for 60 min, and incubated with primary antibodies at their optimized dilutions (supplemental Table 1) – α-cHH3 and α-S100A9 overnight at 4 °C. The tissues were then washed with 3x PBS before being incubated with respective secondary antibodies at room temperature for 1 h. After washing, the tissues were stored in PBS. Whole-mount confocal images were collected using a Leica SP8 microscope.

*In situ* omentum images were also taken using a Leica M205 FA fluorescence stereo microscope. Briefly, shortly following euthanization, mice peritoneal cavities were exposed, and the omentum was located. Because several organs autofluoresce, such as the intestines (Figure S2D), the omentum was gently stretched over the liver, which does not autofluoresce, prior to imaging to minimize background.

### qRT-PCR

The femur and tibia were harvested from 8-week-old C57BL6 mouse. Two ends of the bones were cut and the bone marrow was flushed with a 26g needle filled with cold sterile 1X PBS through a 40μm cell strainer. 10 mL total PBS was used for all four bones. The bone marrow was centrifuged at 500g for 5 minutes at 4°C and then the pellet was resuspended in 500 uL of FACS buffer.

Monocytes were isolated using EasySep^TM^ Mouse Monocyte Isolation Kit (Stem Cell Technologies, #19861) and plated at a density of 200,000 cells per well of a 24 well plate (4 ×10^5^ cells/mL). Monocytes were differentiated in the presence of 20 ng/mL of mCSF in RPMI with 10% FBS and 1x penicillin/streptomycin. After 24 hours in culture, 10ug/mL β-glucan and 33ng/mL IFNγ were added for an additional 48 hours. Total RNA was isolated from cultured cells using TRIzol^TM^ reagent (Invitrogen), according to the manufacturer’s protocols. Glycoblue (Invitrogen) was added as a co-precipitant when handling < 10^6^ cells.

cDNA was synthesized using a High-Capacity RNA-to-cDNA Kit^TM^ (Applied Biosystems #4387406), according to the manufacturer’s protocols. qRT-PCR was performed using SYBR Green PCR Master Mix (Applied Biosystems #4344463) on a QuantStudio^TM^ 3 Real-Time PCR Instrument (Applied Biosystems). The following primers were used for qRT-PCR:

> Vcam1 forward primer 5’-AGTTGGGGATTCGGTTGTTCT-3’;
>
> Vcam1 reverse primer 5’-CCCCTCATTCCTTACCACCC-3’;
>
> C1qa forward primer 5’-AAAGGCAATCCAGGCAATATCA-3’;
>
> C1qa reverse primer 5’-TGGTTCTGGTATGGACTCTCC-3’;
>
> H2-eb1 forward primer 5’-GCGGAGAGTTGAGCCTACG-3’;
>
> H2-eb1 reverse primer 5’-CCAGGAGGTTGTGGTGTTCC-3’;
>
> Ccl8 forward primer 5’-TCTACGCAGTGCTTCTTTGCC-3’;
>
> Ccl8 reverse primer 5’-AAGGGGGATCTTCAGCTTTAGTA-3’;
>
> Il27 forward primer 5’-CTGTTGCTGCTACCCTTGCTT-3’;
>
> Il27 reverse primer 5’-CACTCCTGGCAATCGAGA-3’;
>
> Gapdh forward primer 5’-AGGTCGGTGTGAACGGATTTG-3’;
>
> Gapdh reverse primer 5’-TGTAGACCATGTAGTTGAGGTCA-3’.
>
> All data were normalized to Gapdh quantified in parallel amplification reactions.

### Single cell sequencing and analysis of mouse tumors

Prior to sequencing, immune cells (minus B cells, CD45^+^CD19^-^) were sorted from omentum tumor samples (3 mice/group) and pooled. Samples were uniquely barcoded using TotalSeq-B mouse hashtag antibodies (BioLegend, San Diego, CA), as per manufacturer’s directions, to allow for sample multiplexing for the 10x Genomics Chromium Controller single cell platform (10x Genomics, Pleasanton, CA). Specifically, 1-2 million cells of each sample were first blocked with TruStain FcX PLUS anti-mouse CD16/32 antibody and then incubated with 0.5ug of various anti-mouse hashtag antibodies carrying unique cell barcodes. One 10x G chip lane was loaded with a pool of 4 uniquely barcoded samples and single cell droplets were generated using the Chromium Next GEM single cell 3’ kit v3.1 (10x Genomics). cDNA synthesis and amplification, library preparation and indexing were done using the 10x Genomics Library Preparation kit (10x Genomics), according to manufacturer’s instructions. Overall library size was determined using the Agilent Bioanalyzer 2100 and the High Sensitivity DNA assay and libraries were quantitated using KAPA real-time PCR. One library consisting of a total of 4 samples were pooled and sequenced on the NextSeq 2000 (Illumina, San Diego, CA) using a P3 100 cycle kit (Illumina), paired end run with the following run parameters: 28 base pair x 8 base pair (index) x 90 base pair.

Pre-processing of the scRNA-seq data was performed using Cell Ranger Suite (pipeline v7.0.0, https://support.10xgenomics.com) with refdata-gex-mm10-2020-A transcriptome as a reference to map reads on the mouse genome (mm10) using STAR (Dobin et al., 2013). Cells with over 5% mitochondrial content were filtered out as were those with less than 200 genes with reads to remove cells with low quality and/or cells that are likely dying. The remaining 13534 cells were used for downstream analysis. Batch effect was not observed and hence not corrected for. Seurat v4 (Hao et al., 2021) was used for cell clustering, marker identification, and visualization. The R package SingleR (Aran et al., 2019) was used to determine initial cell types of the clusters using the MouseRNASeq dataset as a reference for cell-specific gene signatures and then verified using known cell-type markers unique to clusters. The MΦ/monocyte clusters were subset and reclustered to identify subclusters of interest. The R package slingshot (Street et al., 2018) was used for trajectory analysis of the MΦ/monocyte subclusters. Differential expression between samples in specific clusters was performed using Wilcoxon Rank Sum Test. Statistically significant differentially expressed genes were used as inputs for enrichment analysis using Qiagen Ingenuity pathway analysis (IPA).

### Single cell RNA sequencing analysis of human OvCa samples

We obtained a single-cell RNA sequencing (scRNA-seq) dataset deposited by Vázquez-García et al from the National Center for Biotechnology Information Gene Expression Omnibus (GSE180661)(Vázquez-García et al., 2022). The dataset consisted of quality-filtered matrices of 929,686 cells. The Seurat package v4.3.0 (RRID:SCR_016341) in R software v4.3.1 (RRID:SCR_001905) and the scanpy package v1.9.3 (RRID:SCR_018139) in Python 3.10 (RRID:SCR_008394) were used for downstream processing. Dimensionality reduction was performed using principal component analysis (PCA) and uniform manifold approximation and projection (UMAP), the same protocol as in the original article. We used the cell type annotation assigned in the original article. The R package scCustomize v1.1.1 and the scanpy function score_genes were used to generate joint plots.

### KM Plotter

The KM Plotter Online tool (https://kmplot.com/analysis/) (Győrffy, 2024) was used to evaluate the relationship between high gene expression and clinical outcome in patients with late stage OvCa (stages III and IV). This open-access TCGA-based database contains bulk RNA sequencing datasets from 1,268 late stage OvCa patients, which allowed us to investigate correlation between overall survival (OS) and enriched genes identified by our scRNA-seq analysis in patients. To analyze the correlation of *IL27/EBI3* coexpression and overall survival, we utilized the multiple-gene analysis and assigned equal weights to *IL27* and *EBI3*.

### In vitro IL27 induction

The femur and tibia were harvested from 8-week-old C57BL6, Syk^MyeΔ^, Dectin-1 KO, or IFNγR-KO mouse. Two ends of the bones were cut, and the bone marrow was flushed with a 26g needle filled with cold sterile 1X PBS through a 40μm cell strainer. 10 mL total PBS was used for all four bones. Bone marrow derived macrophages (BMDMs) were differentiated from total bone marrow cells by growing them in the presence of 20 ng/mL of mCSF in RPMI with 10% FBS and 1x penicillin/streptomycin. Media was supplemented with ½ the volume of initial media+mCSF and differentiation was complete by day 6. After differentiation, the media was changed and BMDMs were stimulated with 10ug/mL β-glucan and 33ng/mL IFNγ for 24-48-hr. IL-27 heterodimer and IL27p28 ELISAs were performed according to their manufacturer protocols (Biolegend, 438707 and R&D Biotechne M2728 respectively).

### T cell activation assay

For T cell activation assay, we treated BMDM with 10ug/mL β-glucan and 33ng/mL IFNγ for 24hr. OT-I T cells (CD8^+^) were isolated from spleens of OT I mice using EasySep™ Mouse CD8^+^ T Cell Isolation Kit (Stem Cell Technologies, 19853A). Dendritic cells (DCs) were isolated from spleens of wild type mice using EasySep™ Mouse Pan-DC Enrichment Kit II (Stem Cell Technologies, 19863). Co-cultures were set up in 4 replicates with BMDM primed as above, OTI T cells, and DCs in the presence of OVA257-264 peptide (Ana Spec Inc., AS-60193-1) at 0.5ng/ml for 3 days. DCs were added at 1:10 ratio to T cells.

For staining surface markers, cells were incubated with fluorescent conjugated antibody cocktail for staining at 4 °C for 30 min. For intracellular staining, cells were stimulated with 20 ng/ml of PMA, 1 μg/ml of ionomycin, 3 μg/ml of brefeldin A, and 2 μM of monensin and incubated at 37 °C for 4 hours. Cells were incubated with antibody cocktail for surface markers at 4 °C for 30 min and then fixed using True nuclear transcription factor buffer set (Biolegend, Cat # 424401) for 20 min at 4 °C. Cell pellets were incubated with antibody cocktail for intracellular markers prepared in permeabilization buffer and incubated at 4 °C for 30 min. All the antibodies were used at 1:400 dilution. Samples were then washed and resuspended in 1x PBS and acquired on BD FACS Symphony flow cytometer. Data were analyzed using FlowJo v10 (Treestar Inc.).

### Statistics

Statistical analyses were performed in Prism (GraphPad Software, Inc.). Statistical tests used and other relevant details are noted in the figure legends. Statistical analysis was performed using the Student’s t test for unpaired samples or one-way ANOVA with a post-hoc Tukey’s multiple comparisons test. Results were considered significant at P < 0.05. Results display all replicated experiments, and presented as mean ± SEM.

## Acknowledgments

We thank Wistar core facilities (imaging, histology, flow cytometry, animal facility, and genomics). We thank Drs. Chris Hunter and Li-Fan Lu for discussions on IL27 and Dr. Maureen Murphy for proofreading the manuscript. This study was supported by NIH Career Enhancement Program through Hopkins-Penn-Wistar Ovarian Cancer SPORE (P50CA228991; N. Zhang), DOD Ovarian Cancer Research Program (OC230051; N. Zhang), W.W. Smith Charitable Trust (C2205; N. Zhang), NIAID (K99AI151198; N. Zhang), NINDS (1R01NS131912; F. Veglia), Cancer Center Support Grant (P30CA010815; N. Zhang), NIH Wistar Training Program in Basic Cancer Research (T32 CA009171; B. Murphy), Japan Society for the Promotion of Science (202360517; T. Miyamoto), and NIH 1R21CA259240 (R. Shinde).

## Author contributions

Conceptualization: N. Zhang, B. Murphy, and T. Miyamoto. Investigation: B. Murphy, T. Miyamoto, B, Manning, T. Kannan, G. Mirji, A. Ugolini, and K. Hamada. Resources: N. Zhang, Y. Nefedova, A. Kossenkov, and R. Zhang. Data interpretation: N. Zhang, B. Murphy, T. Miyamoto, R. Shinde, F. Veglia, L. Huang, D. Claiborne, and Y.P. Zhu. Writing: B. Murphy, T. Miyamoto, and N. Zhang. Supervision: N. Zhang. All authors edited the manuscript.

---

**Supplementary Figure 1.**
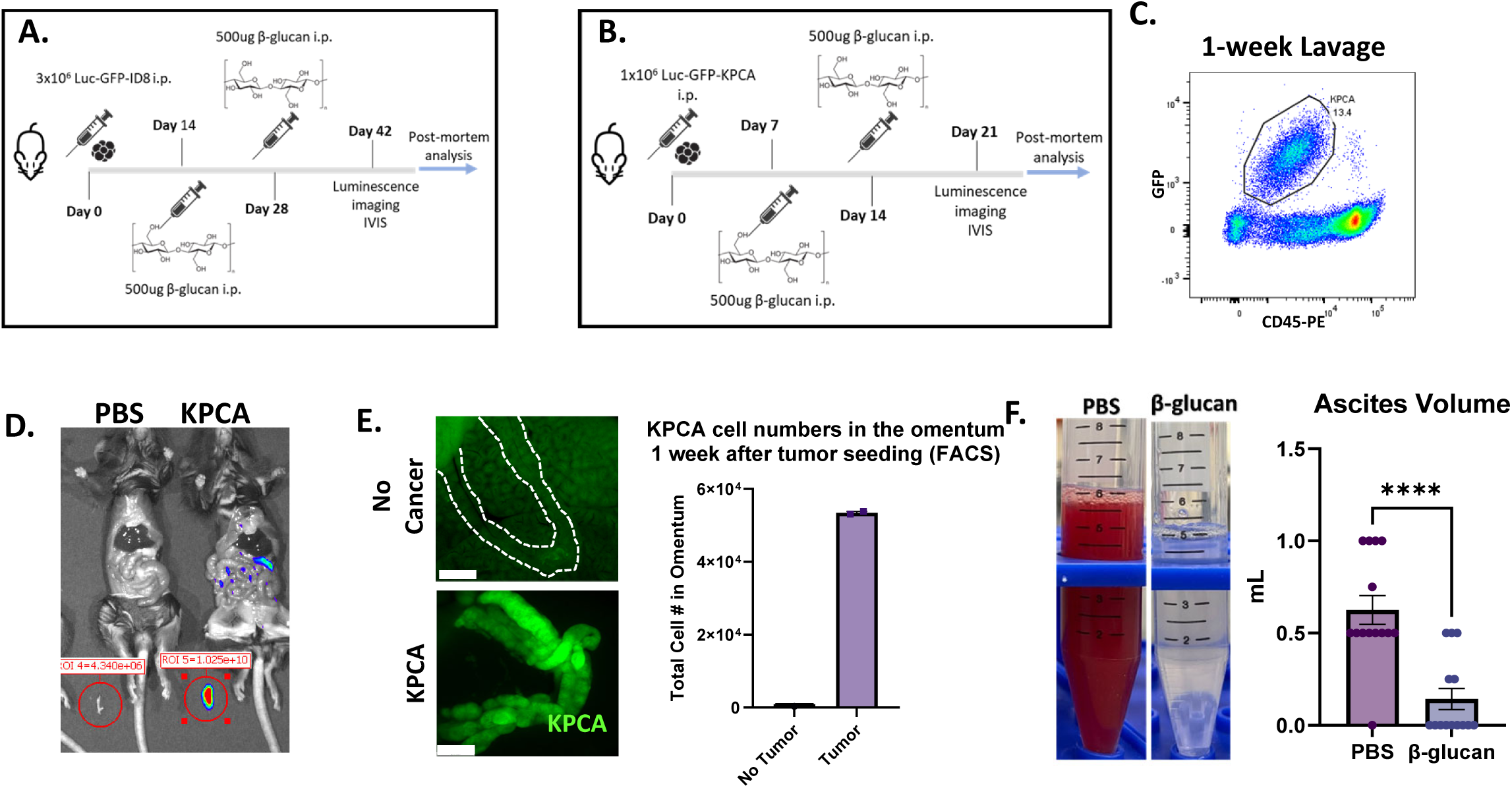
Treatment timelines of (A) ID8 and (B) KPCA tumors treated with β-glucan. (C) Representative flow plot identifying GFP+CD45-KPCA cells in the peritoneal lavage of mice 1 week after tumor seeding. (D) Representative image and quantification of compartmental bioluminescent imaging. The omentum is removed from the cavity; signals (red circle) are obtained separately from the rest of the peritoneal cavity (non-omentum signal). (E) Representative fluorescent images of the omentum and KPCA numbers in the omentum of mice 1 week after KPCA cell injection. Scale bar is 2.5 mm. (F) Representative images of ascites and calculated changes in ascites volumes from PBS- or β-glucan-treated mice. student’s t test was used. ****p<0.0001. Error bars are standard errors of the mean.

**Supplementary Figure 2.**
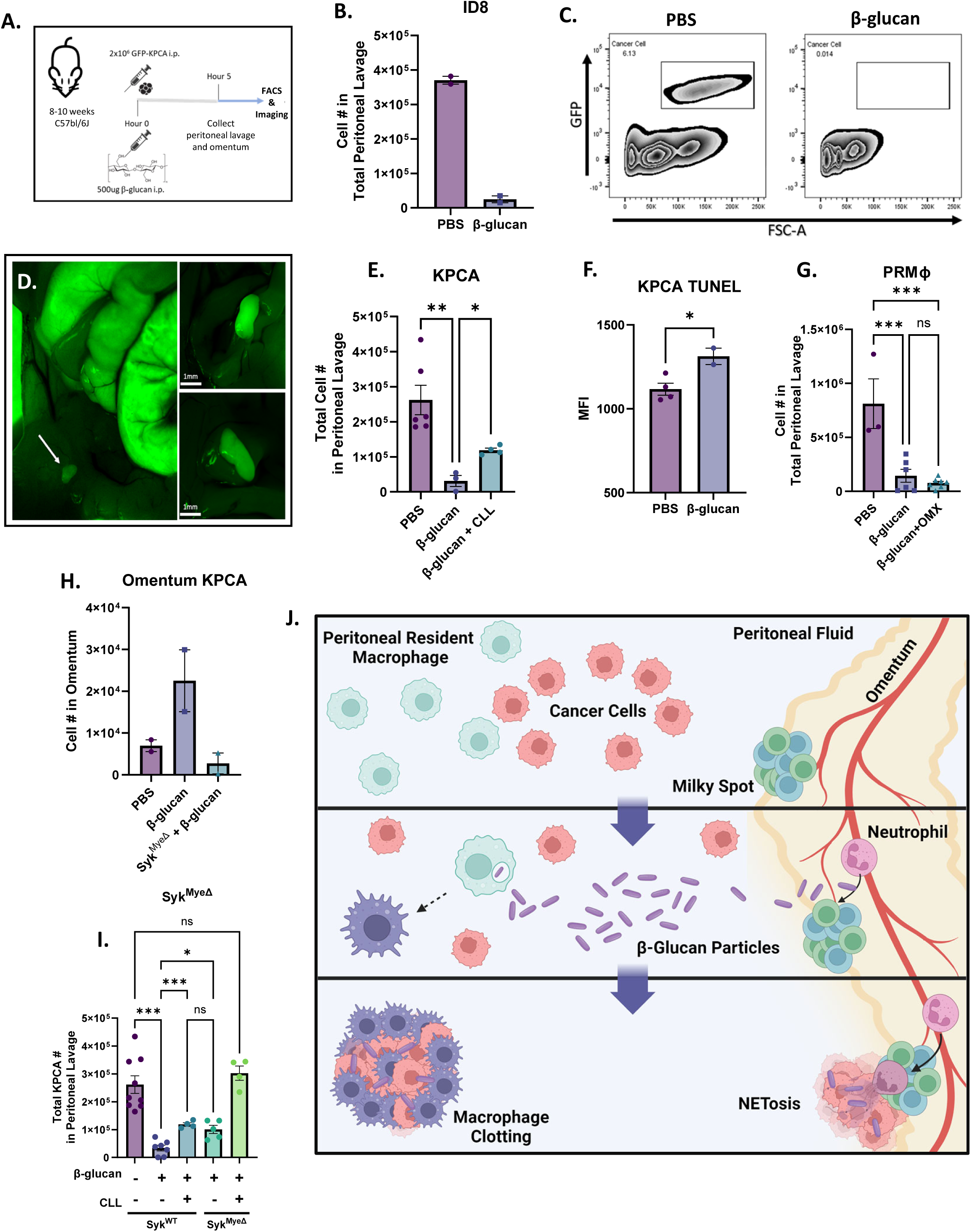
(A) Acute cancer cell capture timeline. (B) Quantification of ID8 cells in the peritoneal lavage of mice 5 hours post β-glucan treatment. (C) Representative flow plots of GFP+ KPCA cells disappearing from the peritoneal lavage 5 hours after intraperitoneal β-glucan administration. (D) Representative *in situ* images of peritoneal clots formed in the peritoneal cavity after β-glucan treatment. These clots contain GFP+ KPCA cancer cells. (E) Quantification of KPCA cells in peritoneal lavage as determined by flow cytometry in control or CLL-pretreated mice 5 hours after intraperitoneal β-glucan administration. (F) MFI of TUNEL staining in KPCA cells in the clots β-glucan treated mice and peritoneal lavage from PBS-treated mice, which do not form clots. (G) Quantification of peritoneal resident macrophages (PRMΦs) in the peritoneal lavage of control or omentectomized (OMX) mice after β-glucan administration. (H) Quantification by flow of GFP+ KPCA cells in the omentum of Syk^WT^ and Syk^MyeΔ^ mice treated with β-glucan. (I) Quantification of KPCA cells in peritoneal lavage in Syk^WT^ and Syk^MyeΔ^ mice 5 hours post indicated treatment with PBS, β-glucan, or CLL. (J) Graphical representation of two mechanisms of cancer cell capture following intraperitoneal injection of β-glucan. One-way ANOVA and student’s t test were used. *p<0.05; **p<0.01; ***p<0.001. Error bars are standard errors of the mean.

**Supplementary Figure 3.**
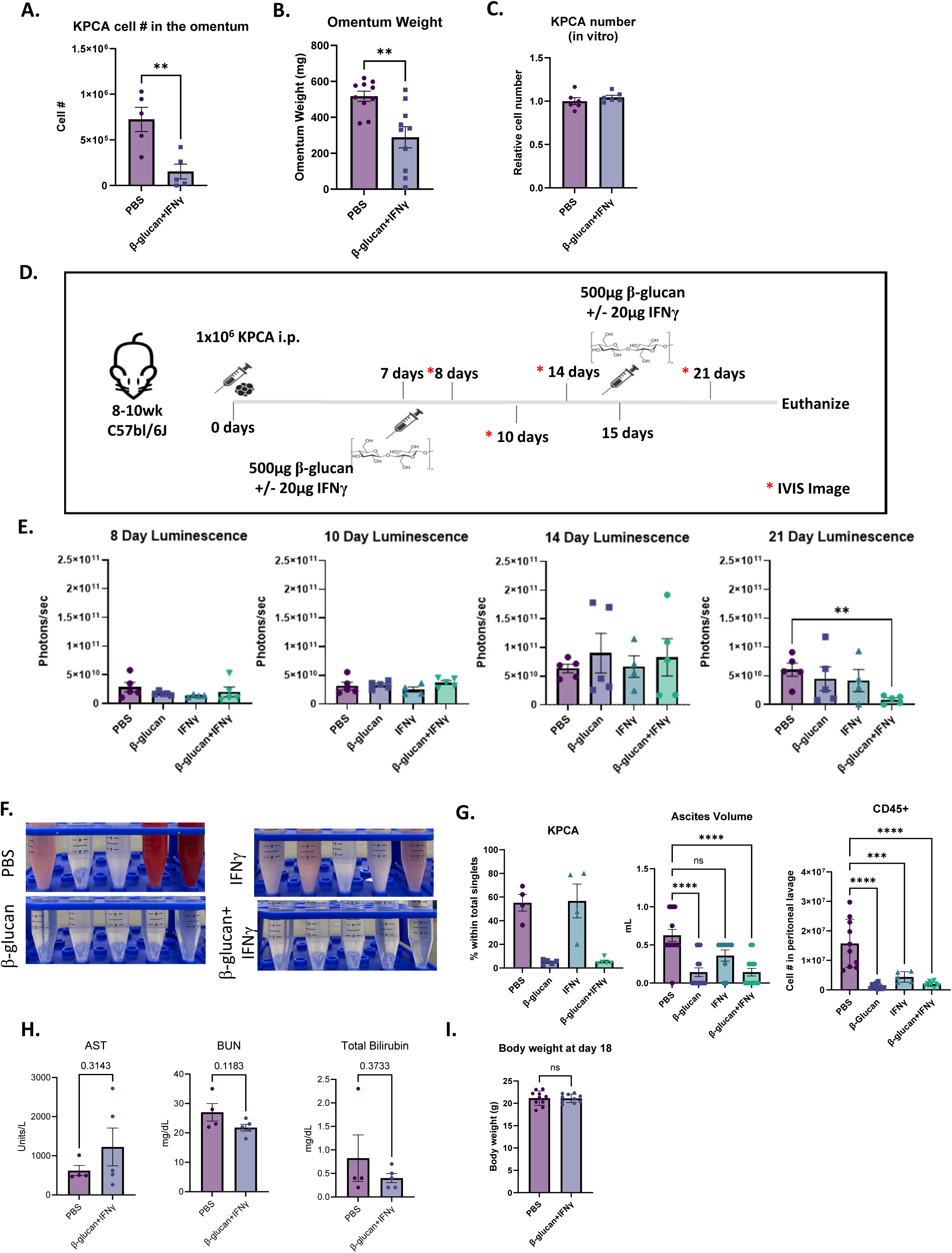
(A) KPCA cell numbers in the omentum evaluated by flow cytometry and (B) Omentum tumor weight in PBS- or BI-treated mice. (C) Quantification of KPCA numbers 48 hours following PBS and BI treatment *in vitro. S*tudent’s t test was used. (D) Treatment and longitudinal imaging timeline in PBS-, IFNγ-, β-glucan-, and BI-treated mice. (E) Quantification of bioluminescence signals in mice tracked longitudinally from day 8 to day 21 after tumor seeding. (F) Representative images of ascites accumulation and (G) quantification of KPCA cells and CD45+ cells and ascites volumes based on the peritoneal lavage of mice treated with PBS, IFNγ, β-glucan, and BI 21 days after tumor seeding. (H) IDEXX clinical chemistry analyses of sera from PBS- or BI-treated mice. (I) Body weight of PBS- or BI-treated mice 18 days after tumor cell seeding. Student’s t test and One-way ANOVA were used **p<0.01, ***p<0.001, ****p<0.0001. Error bars are standard errors of the mean.

**Supplementary Figure 4.**
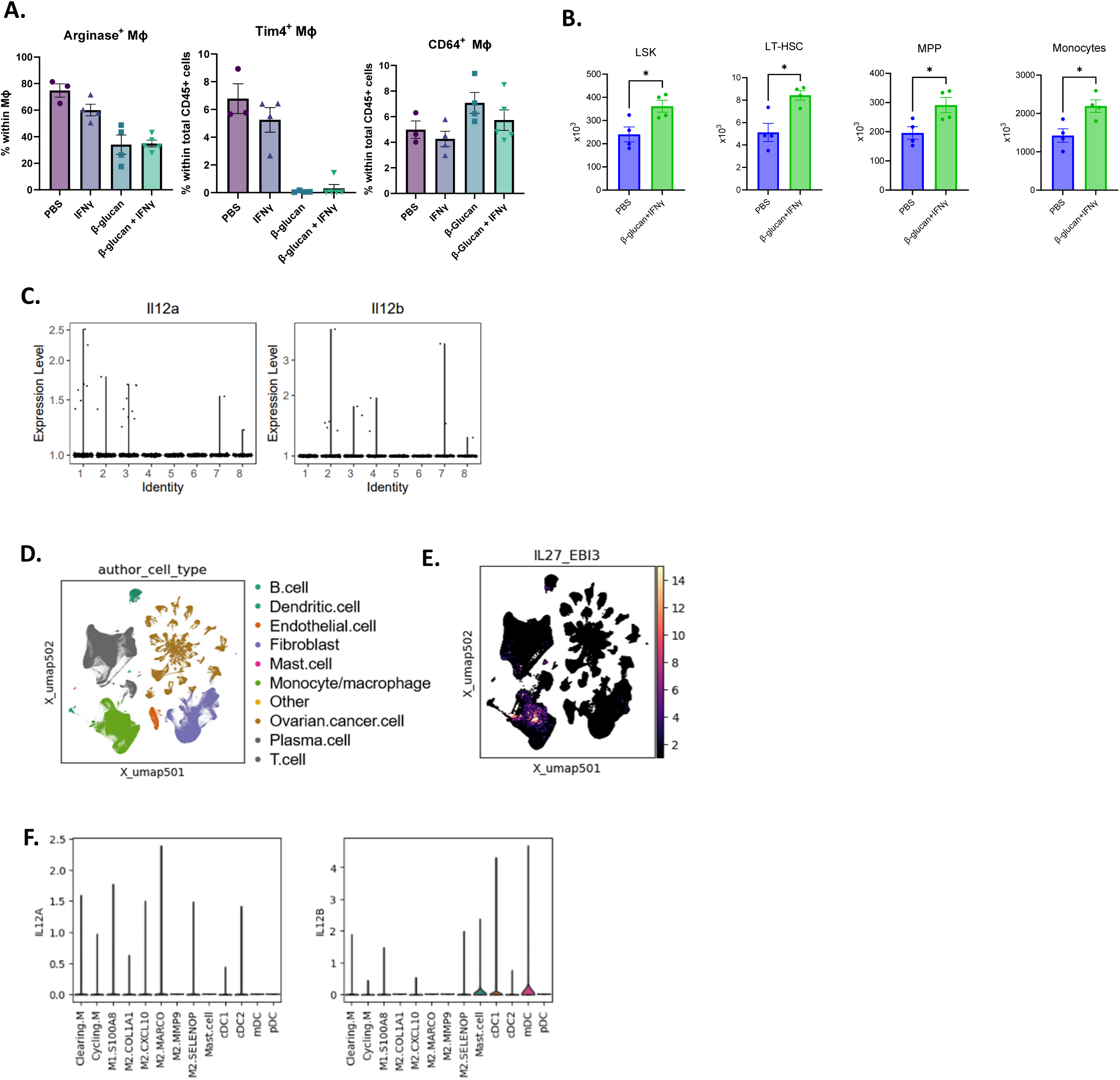
(A) Quantification of frequencies of Arginase+ MΦs, Tim4+ MΦs, and CD64+ MΦs in omentum tumors treated as indicated and determined by flow cytometry. (B) Number of progenitor cells and monocytes in the bone marrow of mice 1 week after PBS or BI treatment. (C) Expression of *Il12a* and *Il12b* in monocyte/MΦ clusters pooed from mice treated with PBS, β-glucan, IFNγ, or BI. (D) UMAP plot of immune cells and co-expression of *IL27-EBI3* in human OvCa patient tumors. (F) Expression of *IL12A* and *IL12B* in each myeloid cell subclusters from human OvCa tumors. Student’s t test was used. *p<0.05. Error bars are standard errors of the mean.

**Supplementary Figure 5.**
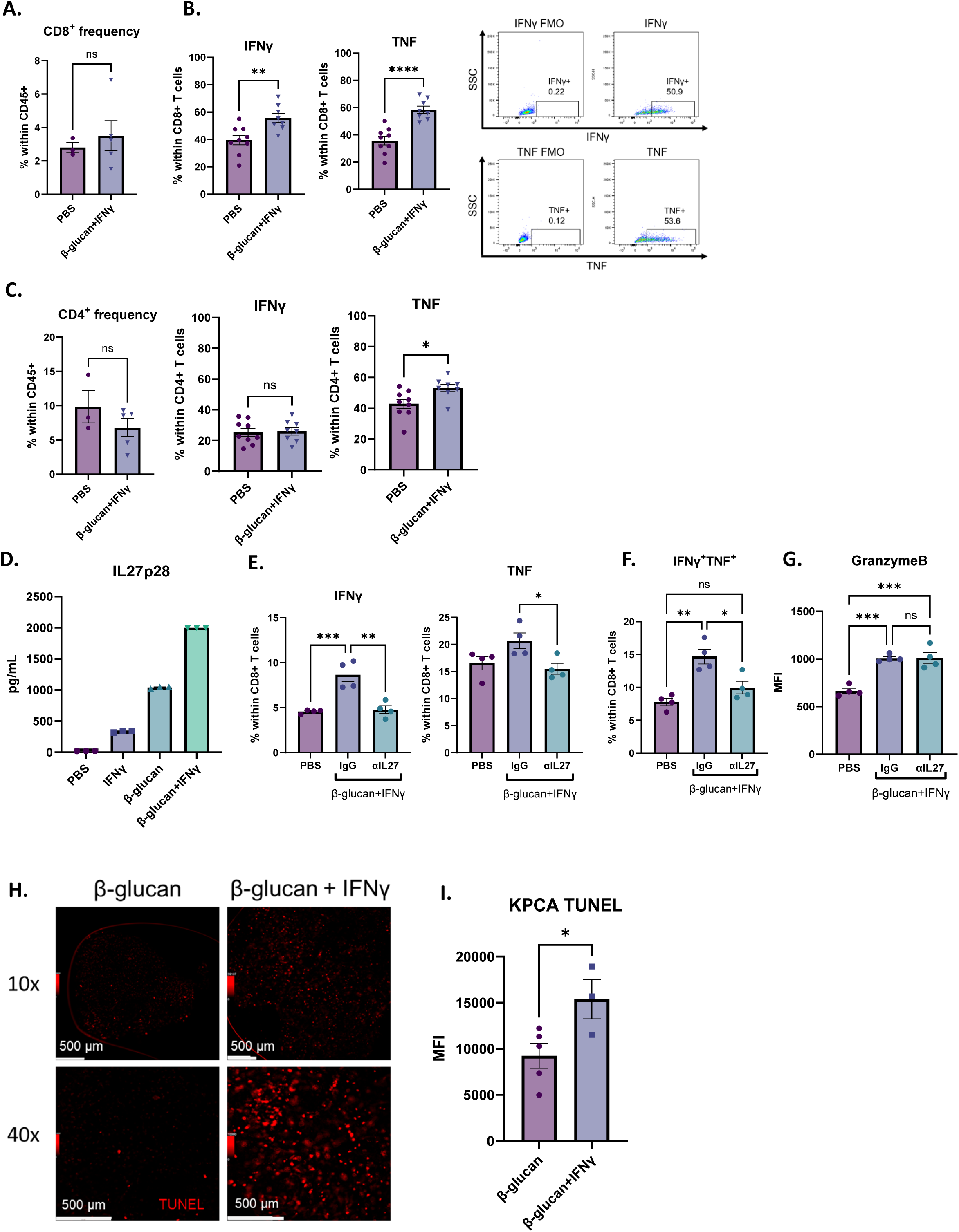
(A) Quantification of frequencies and (B) activation of CD8^+^ T cells in omentum tumors from PBS- or BI-treated mic and flow cytometry plots of TNF- or IFNγ-stained samples, including fluorescence minus one (FMO) plots used to identify positive populations. (C) Quantification of frequencies and activation of CD4^+^ T cells in omentum tumors from PBS- or BI-treated mice. (D) ELISA quantification of IL30 (IL27p28) in supernatant from BMDM cultured with PBS, IFNγ, β-glucan and BI. Frequencies of (E) IFNγ+ or TNF+, (F) IFNγ+TNF+ CD8^+^ T cells, and (G) Granzyme B MFI of CD8^+^ T cells cocultured with MΦ pretreated with PBS or BI in the presence of αIL27 antibody or control IgG. (H) Representative TUNEL staining in β-glucan- or BI-induced clots. (I) FACS quantification TUNEL MFI in GFP+ KPCA Cells. Student’s t test and One-way ANOVA were used. *p<0.05; **p<0.01; ***p<0.001; ****p<0.0001. Error bars are standard errors of the mean.

